# Herpes simplex virus 1 entry glycoproteins form stable complexes prior to and during membrane fusion

**DOI:** 10.1101/2022.05.07.491029

**Authors:** Zemplen Pataki, Andrea Rebolledo Viveros, Ekaterina E. Heldwein

## Abstract

Herpesviruses – ubiquitous pathogens that cause persistent infections – have some of the most complex cell entry mechanisms. Entry of the prototypical herpes simplex virus 1 (HSV-1) requires coordinated efforts of 4 glycoproteins, gB, gD, gH, and gL. The current model posits that the glycoproteins do not interact prior to receptor engagement and that binding of gD to its receptor causes a “cascade” of sequential pairwise interactions, first activating the gH/gL complex and subsequently activating gB, the viral fusogen. But how these glycoproteins interact remains unresolved. Here, using a quantitative split-luciferase approach, we show that pairwise HSV-1 glycoprotein complexes form prior to fusion, remain stable throughout fusion, and do not depend on the presence of the cellular receptor. Based on our findings, we propose a revised “conformational cascade” model of HSV-1 entry. We hypothesize that all 4 glycoproteins assemble into a complex prior to fusion, with gH/gL positioned between gD and gB. Once gD binds to a cognate receptor, the proximity of the glycoproteins within this complex allows for efficient transmission of the activating signal from the receptor-activated gD to gH/gL to gB through sequential conformational changes, ultimately triggering the fusogenic refolding of gB. Our results also highlight previously unappreciated contributions of the transmembrane and cytoplasmic domains to glycoprotein interactions and fusion. Similar principles could be at play in other multicomponent viral entry systems, and the split luciferase approach used here is a powerful tool for investigating protein-protein interactions in these and a variety of other systems.

**IMPORTANCE:** Herpes simplex virus 1 (HSV-1) infects the majority of humans for life and can cause diseases ranging from painful sores to deadly brain inflammation. No vaccines or curative treatments currently exist. HSV-1 infection of target cells requires coordinated efforts of four viral glycoproteins. But how these glycoproteins interact remains unclear. Using a quantitative protein interaction assay, we found that HSV-1 glycoproteins form stable, receptor-independent complexes. We propose that the 4 proteins form a complex, which could facilitate transmission of the entry-triggering signal from the receptor-binding component to the membrane fusogen component through sequential conformational changes. Similar principles could be applicable across other multicomponent protein systems. A revised model of HSV-1 entry could facilitate the development of therapeutics targeting this process.

## INTRODUCTION

Enveloped viruses enter cells by fusing their membrane envelope with a cellular membrane, and most use a single viral protein, termed a fusogen, to perform this function. Fusogens bridge apposing membranes and merge them by refolding from the high energy prefusion conformation into a lower energy postfusion conformation. The energy released during these favorable conformational rearrangements is thought to overcome the activation energy of the membrane fusion process (reviewed in [1]). Fusogens are commonly activated, or triggered, by either exposure to low pH or by binding to a receptor on the target cell (reviewed in [1]). In herpesviruses, the fusion mechanism is more complex, however, and requires three or more viral proteins that join forces to bring about fusion (reviewed in [2]). These double-stranded DNA viruses are significant pathogens that establish lifelong infections (reviewed in [3]). Herpes simplex virus 1 (HSV-1), the focus of this study, is a prototypical herpesvirus (reviewed in [4]) that infects ∼67% of people under the age of 50 worldwide [5] and causes ailments ranging from oral sores (reviewed in [6]) to encephalitis (reviewed in ([7-9]). As there is no curative treatment [10] or vaccines for HSV-1 [11], a better understanding of HSV-1 biology is essential for combating the global burden of HSV-1 disease.

Glycoprotein B (gB) is the conserved fusogen in herpesviruses. By analogy with other fusogens, this homotrimeric protein is thought to merge membranes by refolding from the prefusion [12, 13] to the postfusion conformation [14-18]. However, gB is not a stand-alone fusogen and must be activated by a complex of two conserved viral glycoproteins, gH and gL. In some cases, such as in Kaposi Sarcoma-associated Herpesvirus (KSHV) [19, 20], Varicella Zoster Virus (VZV) [21], or during Epstein-Barr Virus (EBV) infection of epithelial cells [22], gH/gL activates gB upon binding to a cognate host cell receptor directly (reviewed in [2, 23]). In other cases, such as HSV, Human Cytomegalovirus (HCMV), or during EBV infection of B cells, gH/gL instead detects the host cell receptor indirectly by binding to an accessory viral protein (reviewed in [2, 23]), e.g., gD in HSV-1 and 2 ([24, 25] and reviewed in [26]), UL128/UL130/UL131A or gO in HCMV [27-29], and gp42 in EBV [30]. How these multiple glycoproteins interact to bring about fusion is a key question that has not yet been fully answered. While structures of gH/gL bound to accessory viral proteins have been determined for EBV [31] and HCMV [32, 33], and the gH/gL-gB complex in HCMV has been visualized at low resolution [34], very little is known about how HSV-1 gD, gH/gL, and gB interact.

HSV-1 gD, gH, and gB are transmembrane proteins that consist of extracellular, or ectodomains, single-spanning transmembrane domains (TMD), and cytoplasmic domains (CTD or CT), whereas gL is a soluble protein that binds the gH ectodomain. Any of these domains could, in principle, mediate mutual interactions. Indeed, purified recombinant HSV-1 gD and gH/gL ectodomains have been reported to bind *in vitro* in a surface plasmon experiment [35, 36]. Pairwise gD-gH/gL, gD-gB, and gH/gL-gB interactions have also been detected in live cells, by C-terminally tagging each interacting partner with a split-fluorescent protein [37, 38]. However, the split-fluorescent protein approach was not used quantitatively in these studies and has yielded some contradictory results, with some reports suggesting that gH/gL-gB interaction requires the presence of gD and receptor [37, 38] and others maintaining that it does not [39]. As a result, two activation models have emerged. One model is that the viral glycoproteins do not interact until gD binds to a receptor, and then gD binds to gH/gL followed by gH/gL binding to gB for sequential activation (**Fig. 1**) [37, 40, 41]. Alternatively, gD, gH/gL, and gB are already bound to one another, and when a receptor binds to gD, activating signals are transmitted from gD to gH/gL to gB [39]. In both models, the sequential activation of gD, gH/gL, and gB likely involves conformational changes of the glycoproteins ([12, 15, 35, 42] and reviewed in [43]).

**Figure 1.**
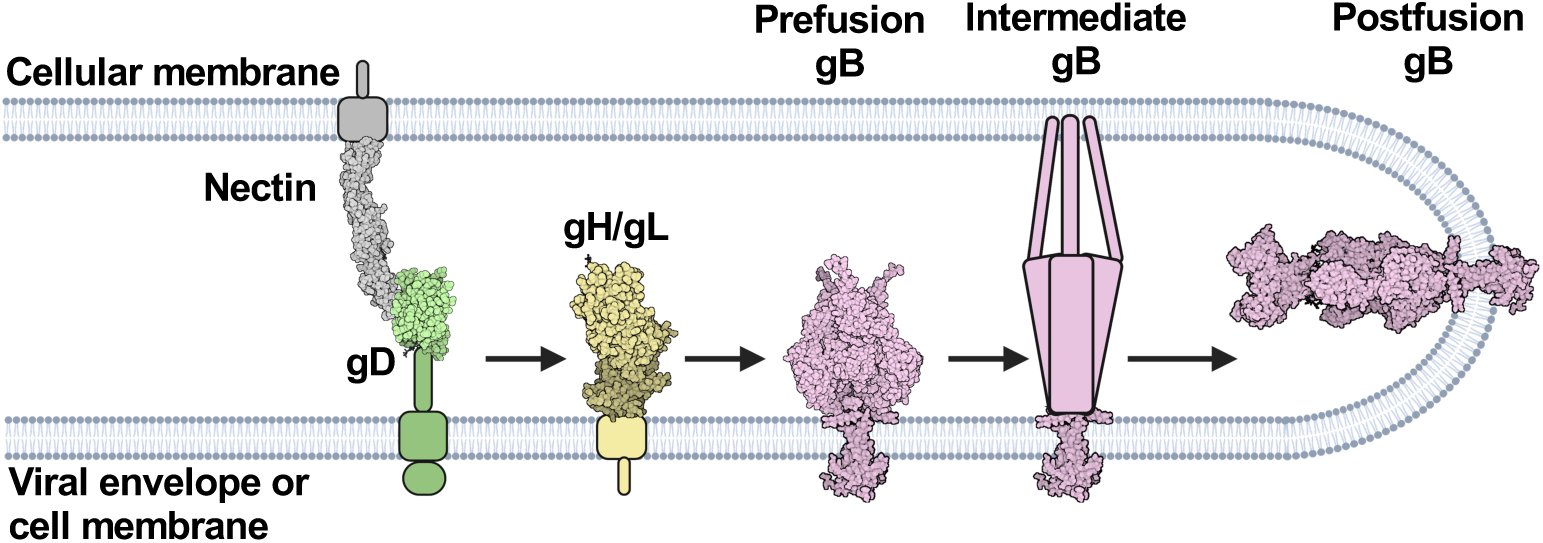
HSV-1 fusion pathway model. gD (2C36) [44] binds a receptor (3U83) [45] on the target cell. gD then activates gH/gL; gH/gL (3M1C) [46] activates gB (6Z9M and 5V2S) [12, 15] to refold and cause membrane fusion. gD is thought to be a dimer [44] but is shown here as a monomer for clarity. Figure created with BioRender.com.

One potential caveat with split fluorescent proteins is that they can generate false positives due to the high-affinity, irreversible interactions of the split fluorescent protein fragments (reviewed in [47]), which may account for the conflicting prior reports. Here, to clarify the timing, dynamics, and sites of interactions between gD, gH/gL, and gB, we turned to a quantitative protein-protein interaction technique called NanoBiT that uses split luciferase fragments fused to putative interaction partners in live cells [48]. Unlike split fluorescent protein fragments, the split luciferase fragments interact in a low-affinity, reversible manner, which reduces the likelihood of false positives and allows the detection of not only complex association but also dissociation over time.

We found that gD-gH/gL, gH/gL-gB, and gD-gB interactions were independent of the presence of the receptor nectin-1 and stable, which suggested that these glycoprotein complexes formed prior to fusion and were maintained throughout fusion. By replacing the HSV-1 gH/gL domains with those of EBV gH or a scrambled sequence, we found that gD-gH/gL and gH/gL-gB interactions involved all three major domains of gH (ectodomain, TMD, and CT). However, while the HSV-1 gH TMD or CT each mediated efficient formation of the gD-gH/gL and gH/gL-gB complexes, the HSV-1 gH ectodomain did not. Therefore, the HSV-1 gH TMD or CT are more important for interactions with gD and gB than the ectodomain. By contrast, all HSV-1 gH/gL domains were essential for fusion suggesting that glycoprotein complex formation is not sufficient for fusion. Finally, our data indicate that whereas gH and gB interact in the endoplasmic reticulum (ER), gH and gD do not.

Putting these findings together, we propose a revised model of HSV-1-mediated fusion whereby a proportion of gH/gL and gB associate in the ER and are transported to the plasma membrane together whereas gD traffics there independently. Once at the plasma membrane, gD, gH/gL, and gB form a complex. Once gD binds to a cognate receptor, the proximity of the glycoproteins within this complex allows for efficient transmission of the activating signal from gD to gH/gL and from gH/gL to gB through conformational changes, in a “conformational cascade”. Our findings increase our understanding of the HSV-1 fusion pathway and may pinpoint new targets for inhibition of HSV-1 infection.

## RESULTS

### HSV-1 gH/gL and gB interact stably and independently of nectin-1

To examine the timing and duration of glycoprotein interactions in HSV-1, we turned to a split-luciferase (NanoBiT) interaction assay in live cells [48]. In this assay, two proteins of interest are tagged with complementary parts of a split luciferase, Lg-BiT and Sm-BiT, and if they interact, the active luciferase is formed, and the resulting luminescence reports on the interaction (**Fig. 2a**). To begin with, gH and gB were C-terminally tagged with Lg-BiT or Sm-BiT (gH-Lg and gB-Sm or gH-Sm and gB-Lg). In initial studies, the gH-Lg/gB-Sm combination resulted in a higher signal-to-noise ratio in the NanoBiT assay than the gH-Sm/gB-Lg combination, so the former combination was chosen for our experiments, as recommended in the NanoBiT technical manual [49]. Both gH-Lg and gB-Sm had a significantly lower total cellular expression than untagged gH and gB (**Fig. 2b-c**), with the Lg tag reducing the expression of gH more than the Sm tag reducing the expression of gB. Both gH-Lg/gL and gB-Sm were expressed on the cell surface, however, indicating that they were properly folded and transported to the cell surface (**Fig. 2d**). The difference in total cellular expression between WT and BiT-tagged constructs was much greater than the difference in their cell surface expression. In other words, the BiT-tagged constructs had relatively high cell surface expression compared to their total cellular expression (**Fig. 2b-d**). By contrast, the WT gH/gL and gB constructs have a relatively low cell surface expression compared to their total cellular expression, which we attribute to protein overexpression. We note that low overall expression levels of gH-Lg and gB-Sm are advantageous because they reduce the potential for non-specific association between interaction partners [49].

**Figure 2:**
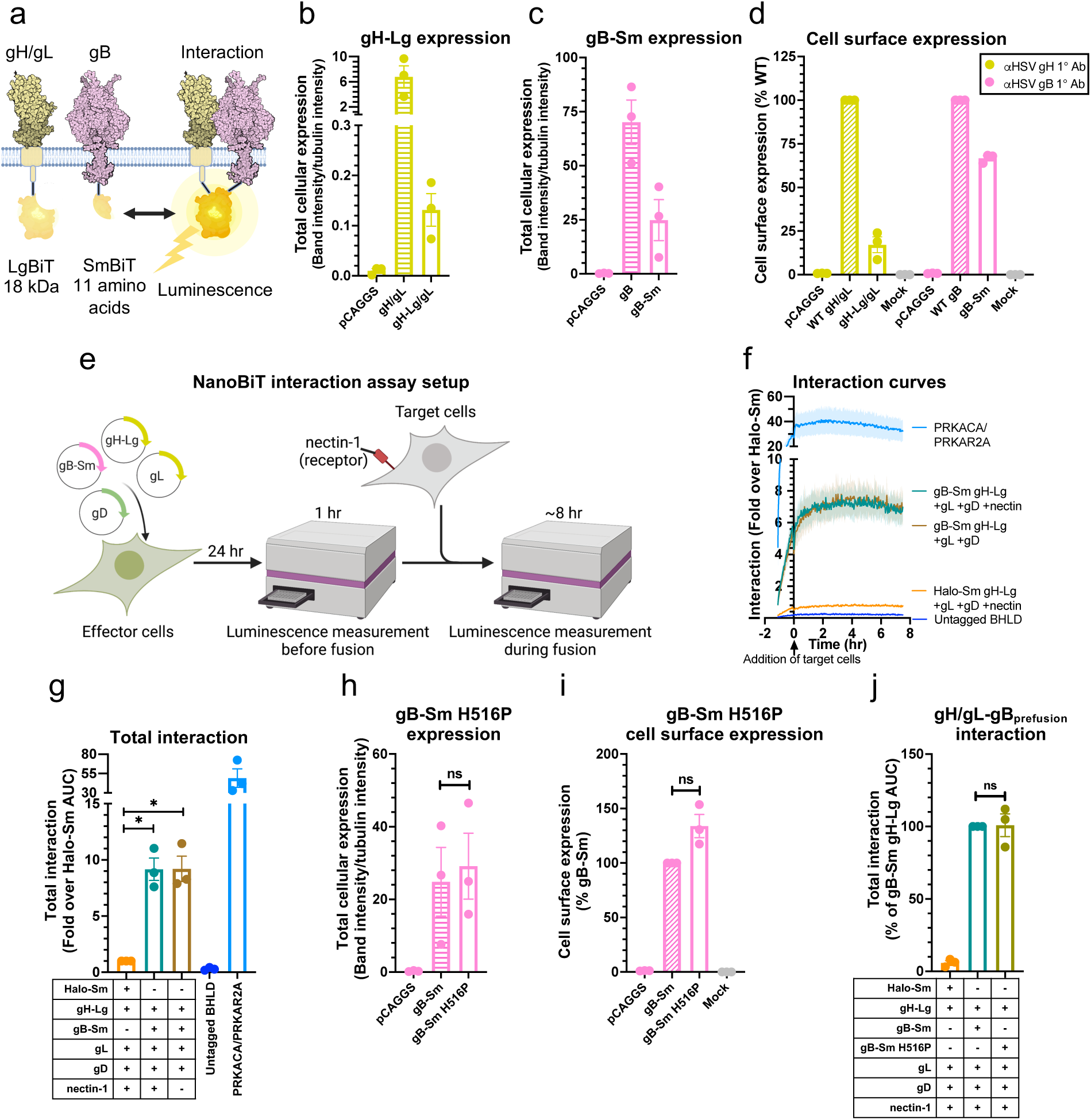
gH/gL and gB interact stably and independently of nectin-1. **a)** NanoBiT protein-protein interaction assay approach [48]. The interaction between gH/gL and gB was tested by tagging gH and gB with complementary parts of a split luciferase and transfecting into cells. Reconstitution of luciferase reports on fusion. **b-d)** Total cellular expression and cell-surface expression of Lg- and Sm-tagged gH and gB by Western blotting and flow cytometry, respectively. R137 and R68 antibodies for gH and gB for Western blotting, respectively. LP11 and R68 antibodies for gH/gL and gB for flow cytometry, respectively. pCAGGS was an empty vector negative control. Mock was an untransfected negative control, incubated with a non-targeting primary antibody. Columns show mean. Error bars are SEM. **e)** Interaction assay experimental setup. Cells are transfected with viral proteins required for fusion, including split-luciferase-tagged proteins of interest. Interaction is measured by luminescence before and during fusion. Fusion is induced by the addition of target cells expressing the viral receptor nectin-1. **f)** gH/gL and gB interaction over time, with target cells expressing or lacking nectin-1. The Halo-Sm and untagged BHLD (gB, gH, gL, gD) conditions were negative controls. PRKACA/PRKAR2A was a positive control. The shaded regions are the SEM. **g)** The interactions are quantified by calculating the area under the curve (AUC). **h-i)** Total cellular expression and cell-surface expression of Sm-tagged gB H516P – which locks gB in its prefusion conformation – by Western blotting and flow cytometry, respectively. **j)** The interaction between gH/gL and gB H516P. Columns show mean. Error bars are SEM. ns: not statistically significant, *: p < 0.05. Data in all panels are three biological replicates from independent experiments. Diagrams were created with BioRender.com.

Interaction between gH/gL and gB was measured prior to and during fusion that was initiated by the addition of target cells expressing the HSV-1 receptor nectin-1 (**Fig. 2e**). Sm-BiT fused to HaloTag (Halo-Sm), which is unlikely to interact with gH, was used as a negative control. HaloTag is derived from a bacterial haloalkane dehalogenase enzyme unrelated to viral glycoproteins (reviewed in [50]). NanoBiT-tagged protein kinase A catalytic (PRKACA) and type 2A regulatory (PRKAR2A) subunits, known to interact, served as a positive control [48]. We found that gH-Lg/gL and gB-Sm interact stably before and during fusion (**Fig. 2f-g**). gH/gL-gB interaction was detected in the absence of the target cells and no noticeable change in gH/gL-gB interaction was seen upon addition of nectin-1-expressing target cells (**Fig. 2f**). Further, the extent of interaction was similar regardless of the presence of nectin-1 in target cells (**Fig. 2g**). We conclude that under these experimental conditions, HSV-1 gH/gL and gB form a stable complex that does not require gD-nectin-1 interaction.

### HSV-1 gH/gL interacts with both prefusion and postfusion forms of HSV-1 gB

Our results suggested that gH/gL and gB interact but left unclear whether gH/gL interacted with the prefusion, postfusion, or both conformations of gB because gB exists as a mixture of these two conformations on the cell surface [51-53] and on virions [54]. Therefore, we introduced the H516P mutation into gB-Sm, which has been reported to stabilize gB in its prefusion conformation [12]. The gB-Sm H516P mutant had similar total cellular expression as the WT gB-Sm (**Fig. 2h**) and was expressed on the cell surface (**Fig. 2i**). The gB-Sm H516P mutant also interacted with gH/gL to a similar extent as the WT gB-Sm (**Fig. 2j**), indicating that gH/gL can interact with prefusion gB, and, therefore, that the gH/gL-gB interaction occurs prior to fusion. Although there is no known gB mutation that stabilizes its postfusion conformation, we hypothesize that gH/gL likely also interacts with postfusion gB because the gH/gL-gB interaction remains stable as fusion progresses, even as the prefusion form converts into postfusion form.

### All HSV-1 gH domains are involved in interactions with HSV-1 gB, but the TMD and the CT are more important than the ectodomain

Having observed a stable gH/gL-gB interaction, we sought to identify the gH domains that were involved in it. Towards this goal, we created gH-Lg variants to disrupt interactions between domains. To disrupt interactions between the ectodomains or TMDs, we replaced the ectodomain or TMD of HSV-1 gH with those of gH from Epstein-Barr virus (EBV) (**Fig. 3a**, constructs 6 and 3, respectively). EBV is a gammaherpesvirus that is distantly related to HSV-1, an alphaherpesvirus, so EBV gH/gL is not expected to interact with HSV-1 gB. Indeed, HSV-1 gB does not mediate cell-cell fusion when paired with EBV gH/gL and vice versa [55]. To enable proper folding, gH chimeras containing HSV-1 gH ectodomain were co-expressed with HSV-1 gL whereas those containing EBV gH ectodomain were co-expressed with EBV gL. To disrupt interactions between the gB CTD and gH CT, the HSV-1 gH CT was scrambled by arranging its amino acids in a random order (**Fig. 3a**, construct 4). This method was chosen instead of replacing the HSV-1 gH CT with that of EBV gH because we reasoned that the scrambled HSV-1 gH CT was less likely to interact with the HSV-1 gB CTD than the EBV gH CT, which shares sequence similarity with the HSV-1 gH CT (**Fig. 3b**). For example, the EBV gH CT has two hydrophobic amino acid clusters, KIV and FFL, that are similar to the KVL and FFW clusters in the HSV-1 gH CT (**Fig. 3b**). Additional constructs were also generated to disrupt interactions in two domains simultaneously (**Fig. 3a**, constructs 5, 7, 8). Finally, the full-length EBV gH-Lg construct was generated as a negative control (**Fig. 3a**, construct 1).

**Figure 3:**
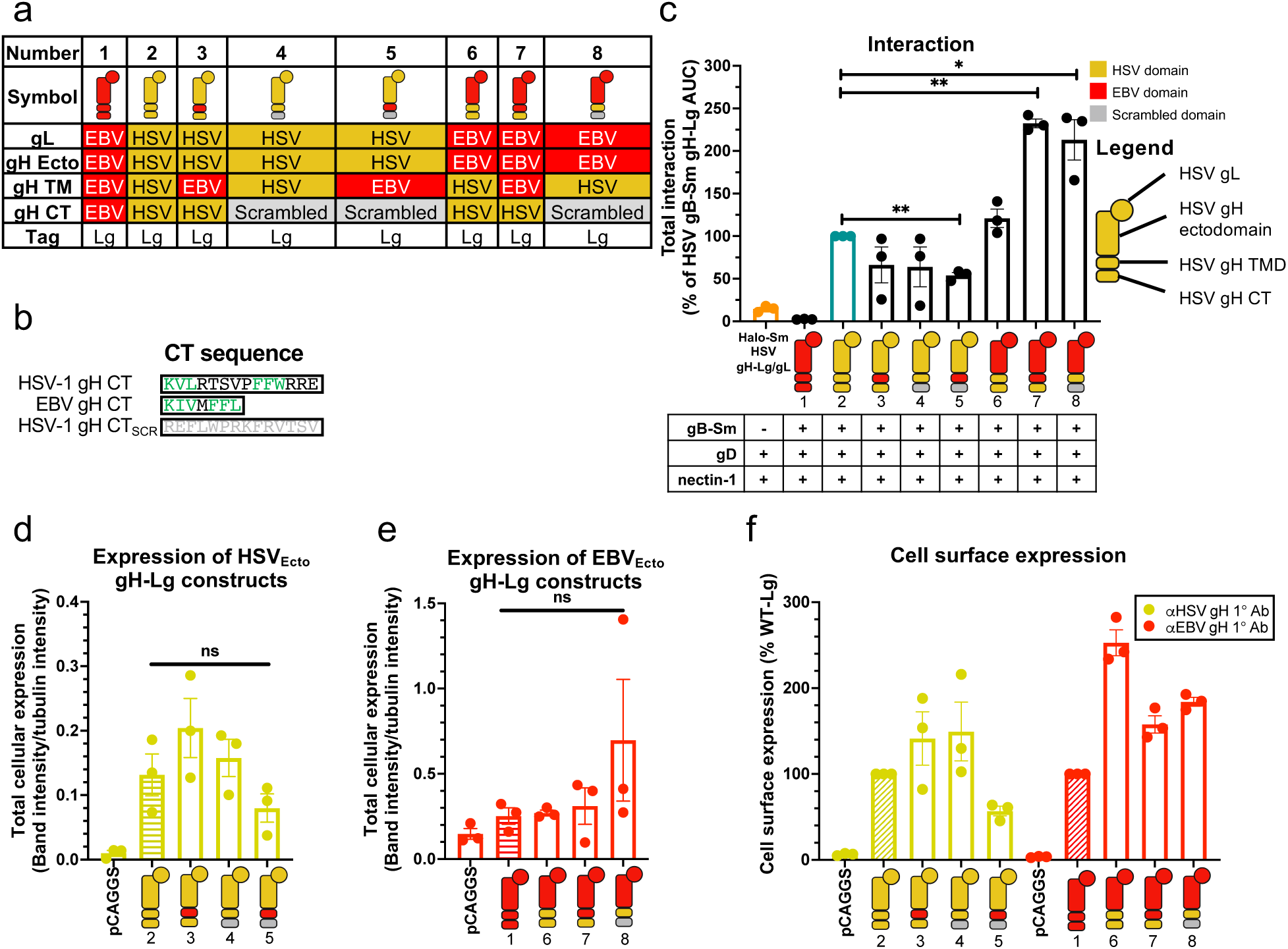
All HSV-1 gH domains are involved in interactions with HSV-1 gB. **a)** Summary of the composition of gH-Lg/gL constructs designed to disrupt domain interactions. **b)** Sequence comparison of HSV-1, EBV, and scrambled HSV-1 gH cytotails. Green indicates similar clusters of residues in HSV-1 and EBV. **c)** Interaction between gB and gH/gL constructs with disrupted ectodomain, TMD, or CTD interactions. Cartoons represent gH-Lg/gL constructs, indicating which domains are HSV-1 (yellow), EBV (red), or scrambled (gray). **d-e)** Total cellular expression of gH-Lg/gL constructs with disrupted domain interactions compared to HSV gH-Lg/gL (2) or EBV gH-Lg/gL (1). R137 and R2267 antibodies were used against constructs with HSV and EBV ectodomains, respectively. **f)** Cell surface expression of gH-Lg/gL constructs with disrupted domain interactions. LP11 and AMMO1 antibodies were used against constructs with HSV and EBV ectodomains, respectively. Columns show mean. Error bars are SEM. *: p < 0.05, **: p < 0.01. Data in all panels are three biological replicates from independent experiments. Cartoons were created using BioRender.com.

As expected, EBV gH-Lg/gL did not interact with HSV-1 gB-Sm (**Fig. 3c**). When single HSV-1 gH domains were replaced, interactions either remained at a WT HSV-1 gH-Lg/gL level (ECTO_EBV_, **Fig. 3c**, construct 6) or were reduced to ∼60% albeit not to a statistically significant extent (TMD_EBV_ and CT_SCR_ constructs, **Fig. 3c**, constructs 3 and 4). We then tested gH constructs in which two domains were replaced simultaneously. When both the gH TMD and CT were replaced (TMD_EBV_-CT_SCR_, **Fig. 3c**, construct 5), interaction decreased to 54%, indicating that when only the gH ectodomain is from HSV-1, gH/gL-gB interaction cannot be maintained at the WT level. Surprisingly, when both the gH ectodomain and TMD, or both the gH ectodomain and CT were replaced simultaneously, interactions increased ∼2-fold (ECTO_EBV_-TMD_EBV_ and ECTO_EBV_-CT_SCR_, **Fig. 3c**, constructs 7 and 8). We conclude that although all gH domains appear to be involved in interactions with gB, when only the TMD or the CT are from HSV-1, they are sufficient to maintain WT-level interactions, whereas the ectodomain is not.

To determine whether differences in interaction were due to differences in the expression levels of the gH-Lg constructs, we measured both the total cellular expression of the constructs by Western blotting as well as their surface expression by flow cytometry. Total cellular expression was measured because gH/gL and gB may interact not only on the cell surface but also at intracellular locations, e.g., in the ER and Golgi. Indeed, it has been shown that gB and gH/gL from cytomegalovirus (CMV), a betaherpesvirus, can interact in the ER [56]. There were no statistically significant differences in total cellular expression of the gH-Lg constructs (**Fig. 3d-e**), so the differences in interaction could not be accounted for by differences in protein expression. All constructs were expressed on the cell surface (**Fig. 3f**), suggesting that they were properly folded.

### All three gH domains are required for fusion

While the TMD or the CT of HSV-1 gH appear sufficient for maintaining WT-level interactions with gB when the rest of the domains are replaced with their EBV gH counterparts or scrambled, it is unlikely that these HSV-1 domains would be sufficient for fusion. To examine this, we tested some of the gH-Lg constructs described above for their ability to support fusion using a split-luciferase cell-cell fusion assay [57]. In this assay, “effector” cells are transfected with gD, gH, gL, gB, and one part of a split luciferase whereas “target” cells are transfected with the nectin-1 receptor and the complementary part of the split luciferase. Effector and target cells are then mixed, and cell-cell fusion is measured by the luminescence produced upon reconstitution of the luciferase (**Fig. 4a**). Fusion of HSV-1 gH-Lg (construct 2) and gB-Sm (in the presence of HSV-1 gD, gL, and nectin-1) was 26% of that of untagged HSV-1 proteins, indicating that the NanoBiT tags do not abrogate the fusion function of gH and gB (**Fig. 4b**). Decreased fusion extent was likely due to the lower cell surface expression of HSV-1 gH-Lg/gL (construct 2, 17% of the untagged HSV-1 gH/gL) and gB-Sm (67% of the untagged HSV-1 gB) (**Fig. 2d**). However, none of the gH-Lg mutant constructs (**Fig. 4b**, constructs 3, 4, 6) were able to support fusion. Thus, interaction between gH/gL and gB is not sufficient for fusion, and all three HSV-1 gH domains are required for fusion.

**Figure 4:**
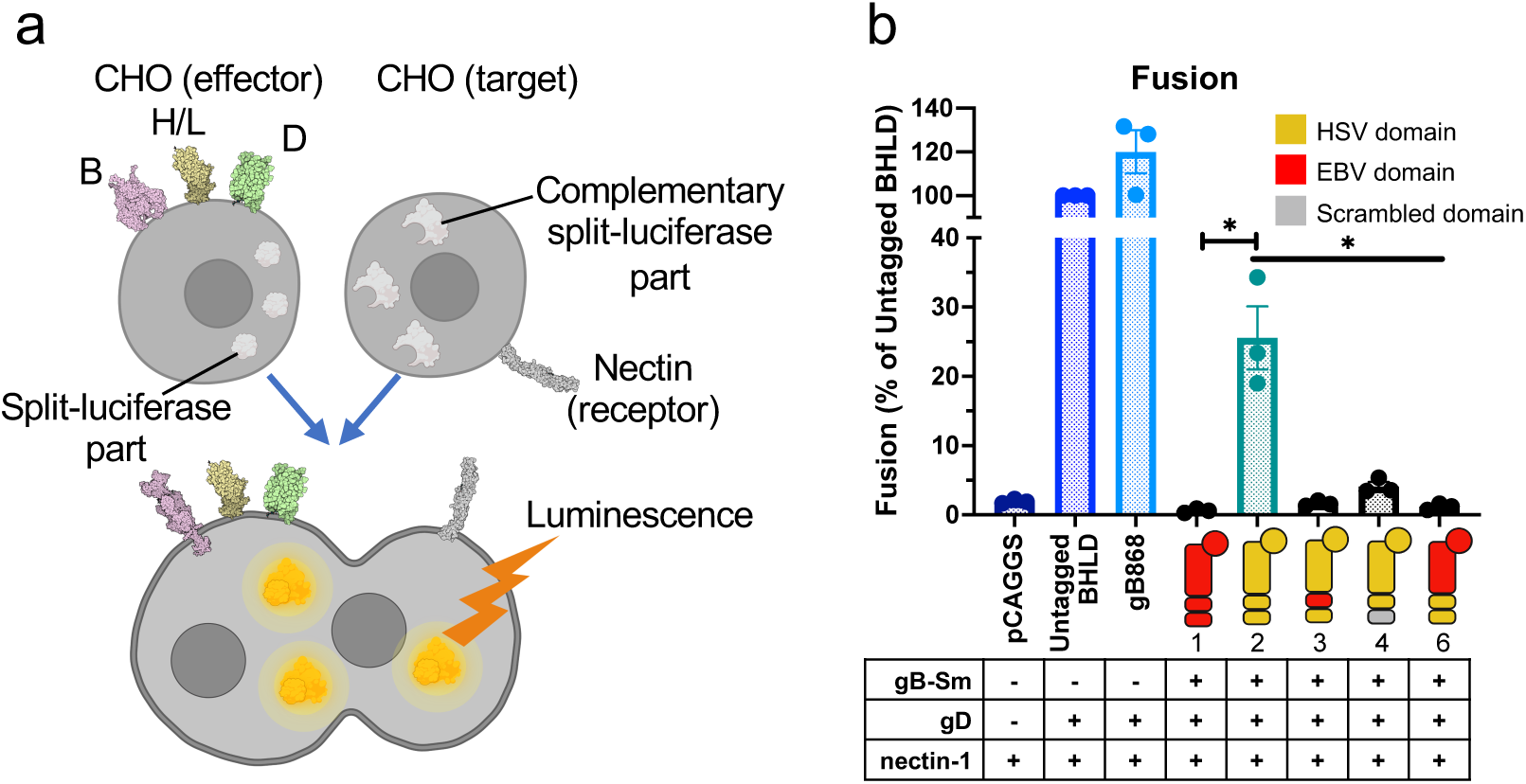
All three gH domains are required for fusion. **a)** Split luciferase cell-cell fusion assay [57] experimental setup. Cells transfected with viral proteins fuse to cells transfected with receptor. Reconstitution of luciferase reports on fusion. **b)** Total fusion of gB-Sm and gH-Lg, including gH-Lg constructs with disrupted domain interactions, 8 hours after mixing effector and target cells. gB868 was a hyperfusogenic positive control gB construct. Columns show mean. Error bars are SEM. *: p < 0.05. Data are three biological replicates from independent experiments. Diagrams and cartoons were created using BioRender.com.

### HSV-1 gH/gL and gD interact stably and independently of nectin-1

The regulatory cascade model of HSV-1-mediated membrane fusion (**Fig. 1**) is predicated upon both gH/gL-gB and gD-gH/gL interactions. To probe the gD-gH/gL interaction and the role of the receptor in it, we generated the gD-Sm construct to test its interaction with gH-Lg/gL (**Fig. 5a**). gD-Sm had reduced total cellular expression (**Fig. 5b**), consistent with other NanoBiT-tagged constructs. gD-Sm was expressed on the cell surface at 68% of the untagged HSV-1 gD (**Fig. 5c**) and supported fusion in combination with gH-Lg at 29% of that of untagged HSV-1 proteins (**Fig. 5d**), indicating that the NanoBiT tag does not abrogate the fusion function of gD. We found that gD-Sm interacted with gH-Lg/gL stably before and during fusion, and that the presence of nectin-1 had no apparent effect on the interaction (**Fig. 5e-f**). We conclude that under these experimental conditions, HSV-1 gD and gH/gL form a stable complex that does not require gD-nectin-1 interaction.

**Figure 5:**
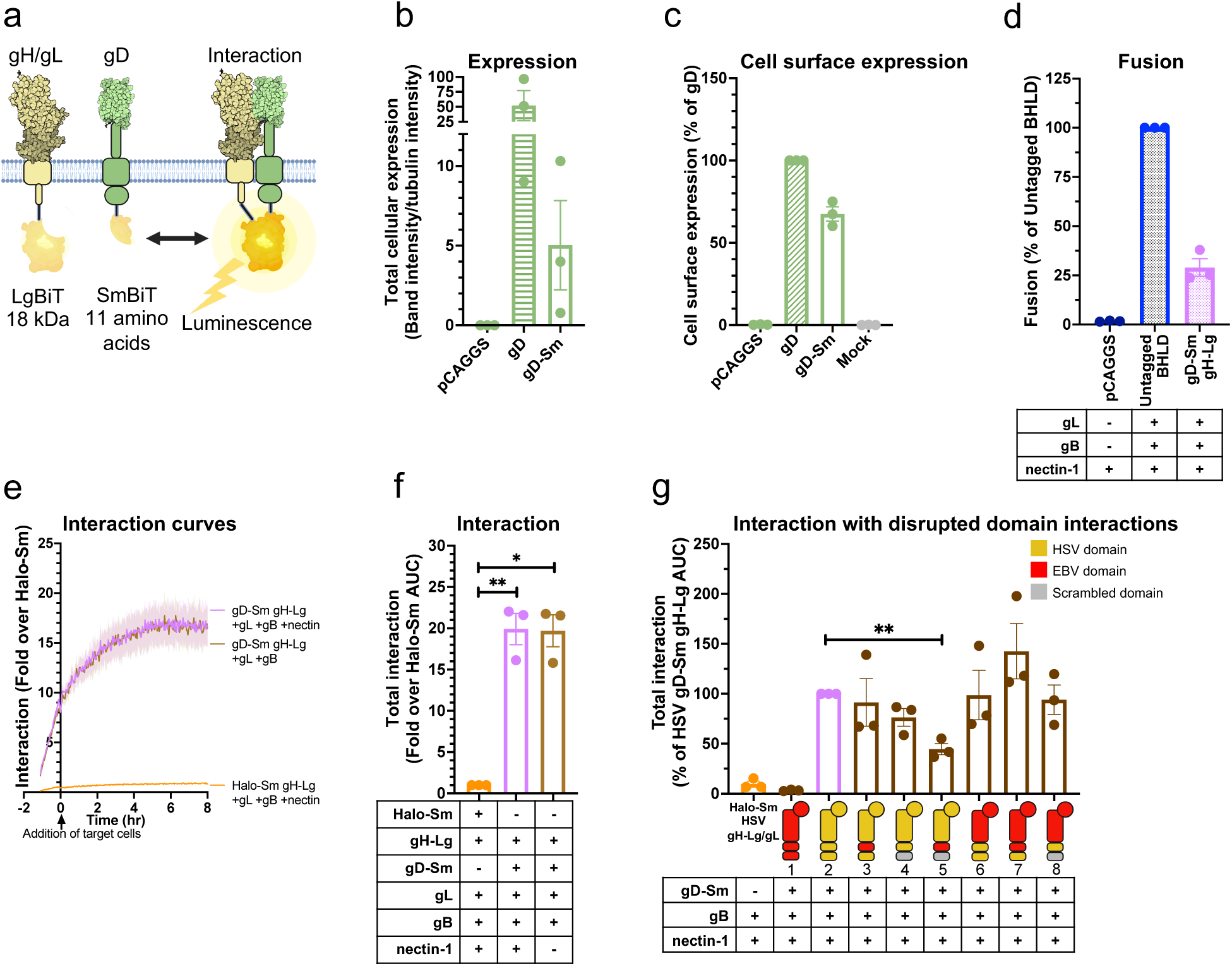
HSV-1 gH/gL and gD interact stably and independently of nectin-1, through all domains. **a)** NanoBiT interaction assay setup to test gD-gH/gL interactions. **b-c)** Total cellular expression and cell surface expression of gD-Sm by Western blotting and flow cytometry, respectively. R7 and DL6 antibodies were used, respectively. **d)** Total fusion of gD-Sm and gH-Lg. **e-f)** Interaction of gD and gH/gL. Curves or bars are the mean and the shaded area or error bars are the SEM. **g)** Interaction of gD with gH/gL with disrupted domain interactions. Columns are mean, error bars are SEM. *: p < 0.05, **: p < 0.01. Data in all panels are three biological replicates from independent experiments in all panels. Illustrations created with BioRender.com.

### All HSV-1 gH domains are involved in interactions with HSV-1 gD, but the TMD and the CT are more important than the ectodomain

To identify gH domains important for the gD-gH/gL interaction, we tested the interaction of gD-Sm with the same series of gH-Lg/gL constructs that were used to probe the gH/gL-gB interaction. As expected, EBV gH-Lg/gL did not interact with HSV-1 gD-Sm (**Fig. 5g**). All but one gH-Lg constructs interacted with HSV-1 gD-Sm at a WT HSV-1 gH-Lg level, with no statistically significant differences. Only the construct in which both the gH TMD and CT were replaced (TMD_EBV_-CT_SCR_, **Fig. 5g**, construct 5) had a significantly decreased interaction, to 45% of WT HSV-1 gH-Lg. Thus, we conclude that although all gH domains are involved in interactions with gD, the TMD or the CT from HSV-1 are sufficient to maintain WT-level interactions whereas the ectodomain is not.

### HSV-1 gD and gB interact stably and independently of nectin-1

Since HSV-1 gH/gL interacted stably with both HSV-1 gB and gD, we asked whether gD and gB could also interact. Thus, we generated HSV-1 gD-Lg to test its interaction with gB-Sm (**Fig. 6a**). Similar to the tagged gH and gB constructs, gD-Lg had low total cellular expression (**Fig. 6b**), was expressed on the cell surface at 12% of the untagged HSV-1 gD (**Fig. 6c**), and supported cell-cell fusion in combination with gB-Sm at 22% of that of untagged HSV-1 proteins (**Fig. 6d**). We found that gD-Lg interacted with gB-Sm stably before and during fusion, and that the presence of nectin-1 had no apparent effect on the interaction (**Fig. 6e-f**). We conclude that under these experimental conditions, HSV-1 gD and gB form a stable complex that does not require gD-nectin-1 interaction.

**Figure 6:**
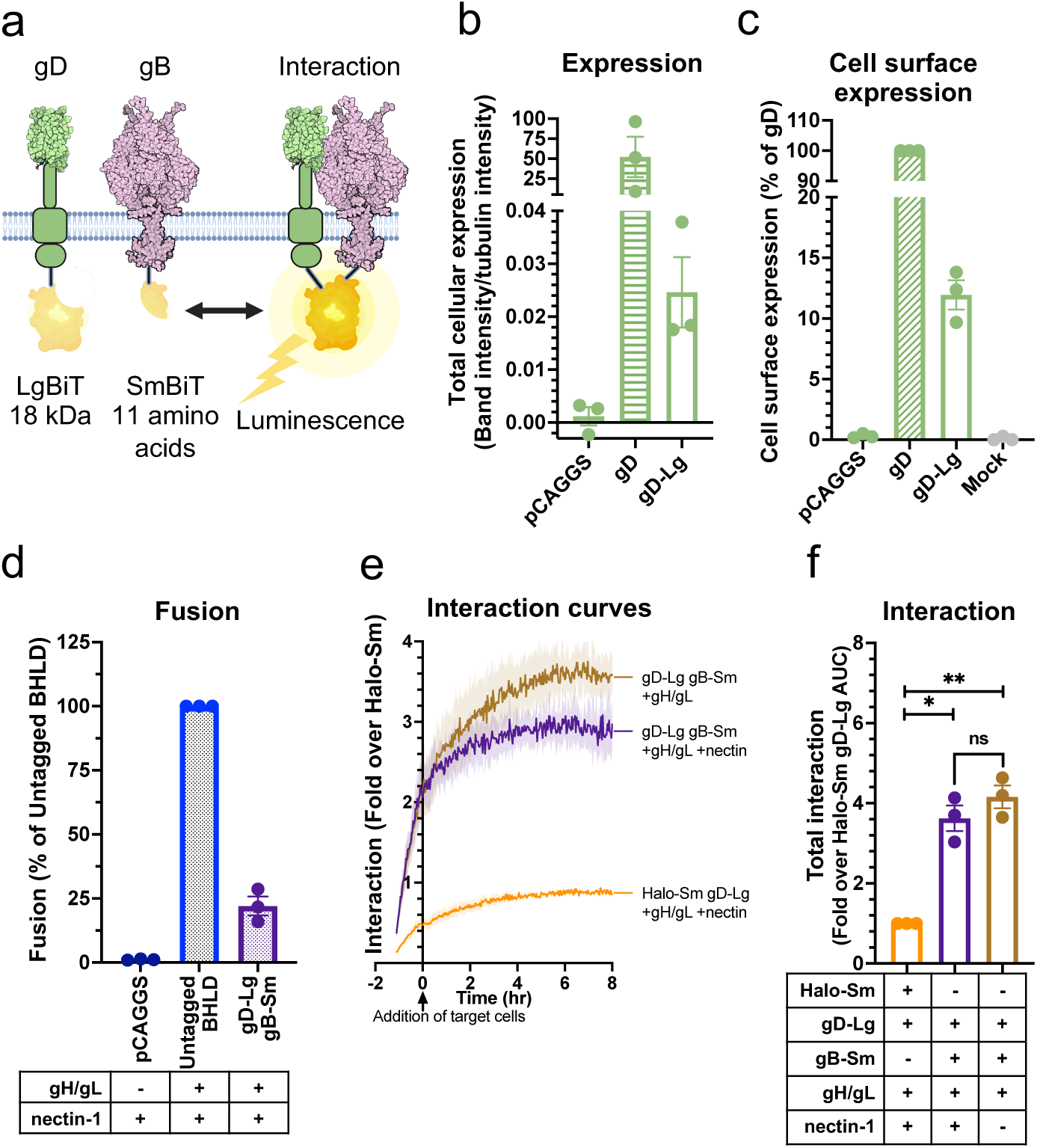
gD and gB interact stably and independently of nectin-1. **a)** NanoBiT interaction assay setup to test gD-gB interactions. Created with BioRender.com. **b-c)** Total cellular expression and cell surface expression of gD-Lg by Western blotting and flow cytometry, respectively. **d)** Total fusion of gD-Lg and gB-Sm. **e-f)** Interaction of gD and gB. Curves indicate mean and the shaded area is SEM. Columns are mean and error bars are SEM. *: p < 0.05, **: p < 0.01. Data in all panels are three biological replicates from independent experiments.

### HSV-1 gD, gH/gL, and gB compete with one another for binding

So far, we have shown that HSV-1 gD, gH/gL, and gB interact with one another in a pairwise manner. However, in all these experiments, all four HSV-1 glycoproteins were present, leaving unclear whether these pairwise interactions required the presence of the third partner. For example, the gD-gB interaction could require the presence of gH/gL. Alternatively, gD, gH/gL, and gB could compete with one another for binding. To differentiate these possibilities, we tested all pairwise interactions in the absence of the third interacting partner. We detected all three complexes, gD-gH/gL, gH/gL-gB, and gD-gB, even in the absence of the third interacting partner (**Fig. 7a-c**). Moreover, we found that all three interactions were reduced in the presence of the third partner, but to a different extent. Whereas gD-gH/gL and gH/gL-gB interactions were minimally reduced in the presence of gB or gD, respectively (**Fig. 7a-b**), the gD-gB interaction was reduced ∼3-fold in the presence of gH/gL (**Fig. 7c**). The observed inhibitory effects are likely due to competition for binding (**Fig. 7d**).

**Figure 7:**
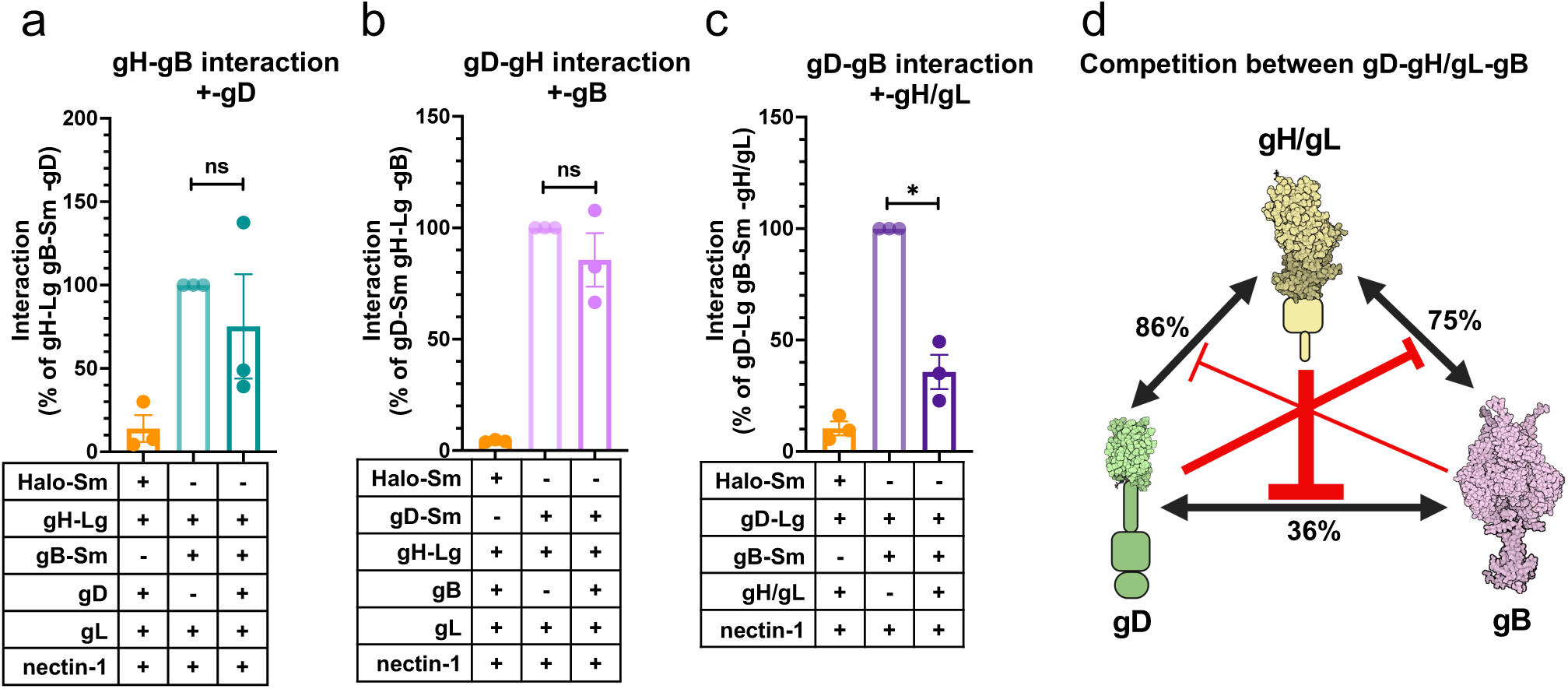
gD, gH/gL, and gB compete with one another for binding. **a)** Interaction of gH and gB in the absence vs the presence of gD. **b)** Interaction of gD and gH/gL in the absence vs the presence of gB. **c)** Interaction of gD and gB in the absence vs the presence of gH/gL. Columns indicate mean and error bars are SEM. *: p < 0.05. Data represent three biological replicates from independent experiments in all panels. **d)** Binding competition model between gD, gH/gL, and gB. The interaction of each pair was decreased in the presence of the third interacting partner, suggesting interaction inhibition by binding competition. The size of the red inhibitory arrows are proportional to the degree of inhibition. Values indicate the % of the interaction of two interacting partners that remains after inhibition by the presence of the third interacting partner. Created using BioRender.com.

### HSV-1 gL is important for gH-gB and gD-gH interactions and for cell surface expression of gH and gB

gL is thought to be required for transport of gH to the cell surface [58]. We confirmed that the total cellular expression levels of gH-Lg were similar in the presence or absence of gL (**Fig. 8a**) and that gH was not expressed on the cell surface without gL (**Fig. 8b**). Next, we examined the location of gD-gH and gH-gB interactions. In the absence of gL, gH localizes to the ER [59]. Therefore, if gD-gH and gH-gB interactions occurred only on the cell surface, they would no longer be detectable in the absence of gL. Conversely, if interactions occurred in the ER, they could be maintained in the absence of gL. We found that, in the absence of HSV-1 gL, the gD-gH interaction reduced nearly to background levels, ∼6-fold (**Fig. 8c**), which suggested that gD and gH/gL interact mainly on the cell surface. In contrast, in the absence of gL, the gH-gB interaction decreased ∼2-fold (**Fig. 8d**), suggesting that gH and gB can interact in the ER at ∼50% of the level of gH/gL-gB interactions under our experimental conditions.

**Figure 8:**
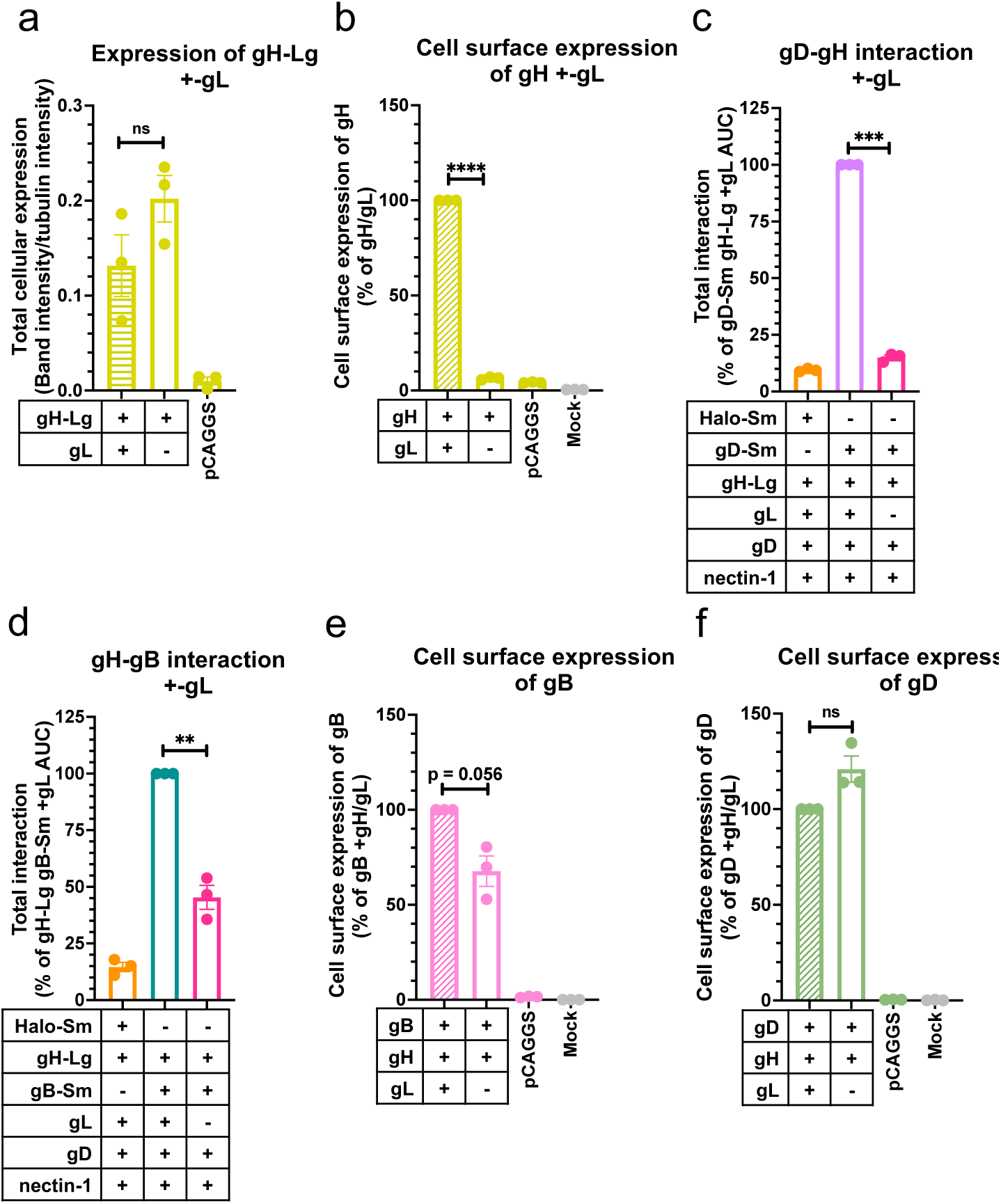
HSV-1 gL is required for gH-gB and gD-gH interactions and for cell surface expression of gH and gB. **a)** Total cellular expression of gH-Lg in the presence vs the absence of gL. **b)** gH cell surface expression in the presence vs the absence of gL [59, 60] using R137. **c)** gD-gH interaction in the presence vs the absence of gL. **d)** gH-gB interaction in the presence vs the absence of gL. **e)** gB cell surface expression in the presence vs absence of gL. **f)** gD cell surface expression in the presence vs absence of gL. Columns are mean. Error bars are SEM. **: p < 0.01, ***: p < 0.001, ****: p < 0.0001. Data represent three biological replicates from independent experiments in all panels.

Since in the absence of gL, gH is unable to leave the ER yet can still interact with gB, we next tested whether the gH-gB interaction can also prevent gB from leaving the ER to traffic to the cell surface. Indeed, the gB cell surface expression decreased to 68% in the presence of gH alone relative to when both gH and gL were present (**Fig. 8e**). In contrast, the gD cell surface expression was not noticeably affected by the absence of gL (**Fig. 8f**). We conclude that gH and gB can interact in the ER and that gH/gL and gB may traffic together to the cell surface, while gH and gD do not interact in the ER to any significant extent and that gH/gL and gD traffic to the surface independently from one another (**Fig. 9a**).

**Figure 9:**
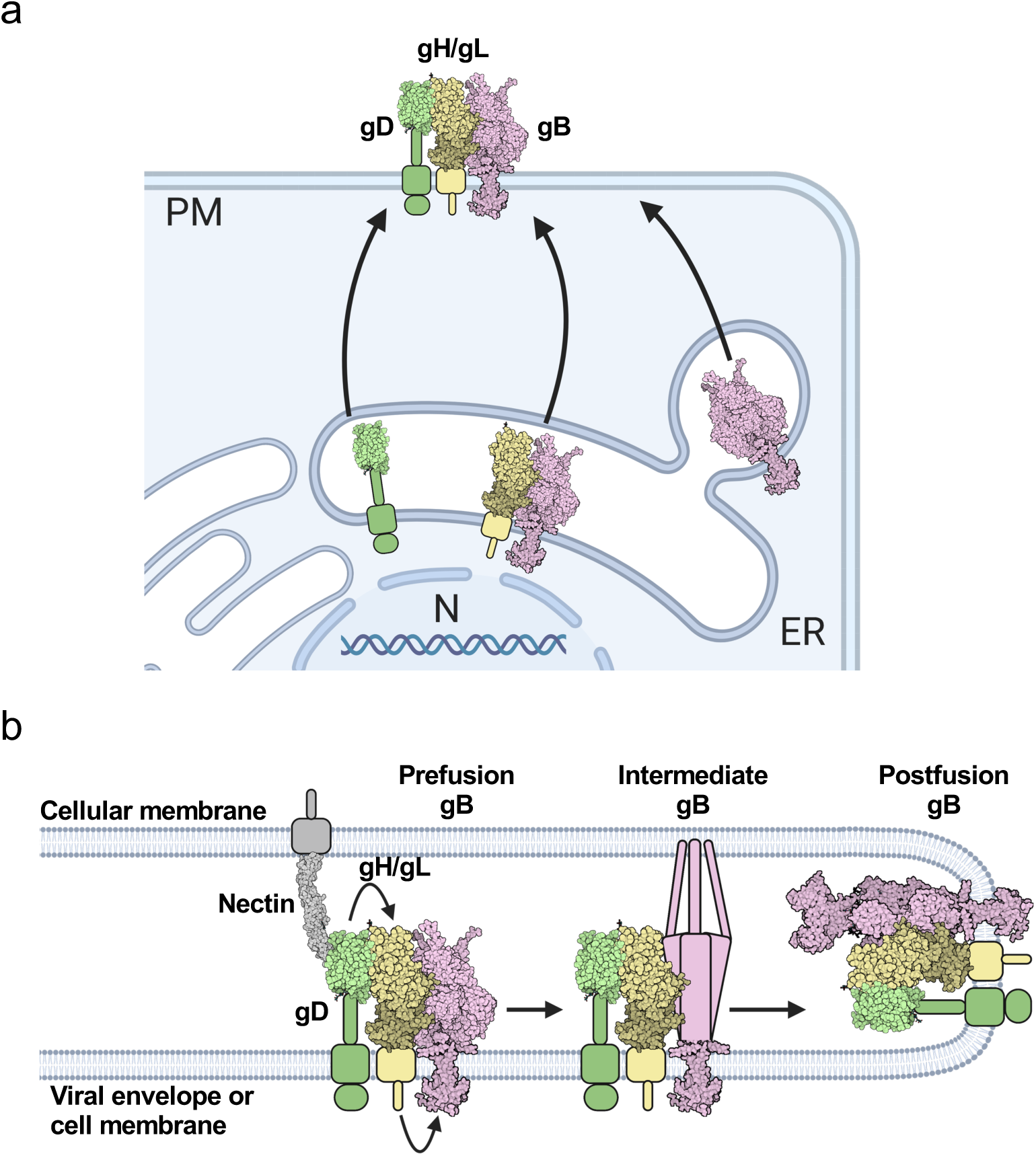
gD-gH/gL-gB trafficking, interaction, and fusion models. **a)** Intracellular interactions and trafficking of gD, gH/gL, and gB. gD does not interact with gH in the ER and traffics independently of gH to the plasma membrane. gB interacts with gH in the ER and may traffic with gH/gL to the plasma membrane. Some gB traffics to the plasma membrane without gH/gL. gD, gH/gL, and gB all interact with one another once they leave the ER and compete with one another for binding. gH/gL inhibits binding of the other two binding partners the most, suggesting it binds well to both gD and gB and may position itself between gD and gB in the putative gD-gH/gL-gB complex. gH/gL interacts with gD and gB through all three domains. Cell and glycoprotein sizes are not to scale. **b)** New HSV-1 fusion pathway model. gD, gH/gL, and gB are all interacting with each other before fusion. Nectin binds to gD, causing a conformational change (not shown), which activates gH/gL via their ectodomains. gH/gL undergoes a conformational change (not shown) and the gHCT activates the gBCTD. The gB ectodomain refolds and catalyzes membrane fusion. gD, gH/gL, and gB continue to interact. For this model we assume a complex with a 1:1:1 ratio of gD:gH/gL:gB, but the true stoichiometry is unknown.

## DISCUSSION

### Interactions between gD, gH/gL, and gB occur prior to and during fusion

Using the NanoBiT interaction assay, which uses split luciferase [48], we established the timing, duration, and dynamics of the interactions between HSV-1 gD, gH/gL, and gB. The advantage of the NanoBiT interaction assay over the commonly used split fluorescent proteins is that the split luciferase fragments interact in a low-affinity, reversible manner. This reduces the likelihood of false positives and allows the detection of not only complex association but also dissociation over time. As far as we know, this study is the first application of the split luciferase approach to probing interactions between membrane proteins. The split luciferase approach is thus a powerful tool that may be broadly applicable across a variety of systems.

We found that pairwise gD-gH/gL, gH/gL-gB, and gD-gB interactions occurred prior to fusion, remained stable throughout fusion, and were independent of the presence of the receptor nectin-1. As an example, gH/gL interacted with gB in the absence of gD and also interacted with the gB H516P mutant that is supposed to lock gB in the prefusion conformation [12]. These findings support the notion that gH/gL-gB interactions occur prior to fusion [39]. The NanoBiT signal for all three pairs, gD-gH/gL, gH/gL-gB, and gD-gB, remained stable upon addition of the receptor-expressing target cells, suggesting that the complexes do not dissociate as gB refolds from the prefusion to the postfusion conformation (**Fig. 9b**). Therefore, we conclude that the gD-gH/gL, gH/gL-gB, and gD-gB complexes form independently of fusion and are maintained throughout the fusion process.

The detection of gD-gH/gL, gH/gL-gB, and gD-gB interactions using the split-luciferase assay is consistent with previous studies that observed these interactions using split fluorescent proteins [37, 38]. However, our observation that the gH/gL-gB interaction occurs prior to fusion, agrees with some reports using split fluorescent proteins [39] but not others [37, 38]. We hypothesize that this could be due to a higher sensitivity of the NanoBiT split-luciferase approach used here for detecting interactions ([48] and reviewed in [47]). Alternatively, gD-receptor binding could increase the rate of gH/gL-gB association/dissociation, which could increase the signal due to the irreversibility of the split-fluorescent protein interactions, explaining previous observations [37].

### gD-gH/gL and gH/gL-gB interactions involve all HSV-1 gH domains, but the TMD and CT are more important than the ectodomain, and all three domains are required for fusion

Replacing domains of the HSV-1 gH with those of EBV gH or a scrambled sequence revealed that all three domains were involved in gD-gH/gL and gH/gL-gB interactions. For example, when the TMD or CT were the only endogenous HSV-1 gH domains, the chimeric gH could maintain WT-level interactions with HSV-1 gD and gB. However, when the ectodomain was the only endogenous HSV-1 gH domain, interaction with HSV-1 gD and gB decreased ∼2-fold. Therefore, gD-gH/gL and gH/gL-gB interactions through their TMDs and CTDs are greater than through their ectodomains. Whereas previous studies focused mainly on ectodomain interactions [35, 36, 61], our study highlights the previously overlooked yet important contributions of the TMD and CTDs to HSV-1 glycoprotein interactions.

Surprisingly, when the TMD or CT were the only endogenous HSV-1 gH domains present, the gH/gL-gB interaction was enhanced ∼2-fold. One possible explanation for this could be a competition for binding of gB to the HSV-1 gH/gL ectodomain by an unknown host protein. Such a competitor would be unable to bind the constructs containing the EBV gH/gL ectodomain, which could explain increased binding of those constructs to HSV-1 gB relative to WT HSV-1 gH/gL. Indeed, integrins have been reported to bind HSV-1 gH/gL [62, 63].

While each of the HSV-1 gH/gL domains, to a certain extent, could maintain interactions with gD and gB, replacing any of the HSV-1 gH/gL domains with those of EBV gH or a scrambled sequence significantly reduced fusion. This suggests that all three HSV-1 gH/gL domains are required for fusion and that they each perform a specific function in fusion beyond interaction with gD and gB, such as transduction of the triggering signal from gD to gB.

### gH/gL blocks gD-gB interactions

We found that the gD-gH/gL, gH/gL-gB, and gD-gB interactions could occur in the absence of the third interacting partner. Moreover, these pairwise interactions were greater when the third partner was absent, which suggested that each interacting partner inhibited the interactions of the other two partners. Since all three glycoproteins interact with one another, this inhibitory effect could be due to competition based on shared or overlapping binding sites, allosteric effects, or steric crowding. Interestingly, the inhibitory effect of gD on the gH/gL-gB interaction and gB on the gD-gH/gL interaction was relatively modest and statistically insignificant. By contrast, gH/gL caused a statistically significant, ∼3-fold reduction in gD-gB interaction. This suggests that gD and gB preferentially bind to gH/gL than to each other, which positions gH/gL in between gD and gB in a putative gD-gH/gL-gB complex and supports the role of gH/gL as the “middleman” between gD and gB (reviewed in [64]).

### gD and gB differ in their trafficking and interactions with gH/gL

HSV-1 gD, gH/gL, and gB are produced in the ER, traffic to the plasma membrane, and become endocytosed, ending up in vesicles derived from the endosomes or Trans Golgi network that serve as virion assembly sites during infection [65, 66]. Alphaherpesvirus glycoprotein traffic from the ER to the plasma membrane is thought to occur by the exocytic pathway [67], but whether HSV-1 gD, gH/gL, and gB interact during this process is unknown. Once at the plasma membrane, HSV-1 gB can endocytose independently of other viral glycoproteins to reach the virion assembly site whereas gH/gL and gD require colocalization – and likely interaction – with other glycoproteins such as gM because they lack an endocytic signal [66].

Given the extensive interactions between gD, gH/gL, and gB that we observed in this study, we asked where in the cell they interact and whether these proteins co-traffic to the cell surface. By taking advantage of the inability of gH to leave the ER without gL, we found that in the absence of gL, gD-gH interactions in the ER were minimal whereas gH-gB interactions in the ER were about 50% of the level of overall gH/gL-gB interactions. This suggests that gB interacts with gH to a far greater extent in the ER than gD does. In addition, we found that when gH was expressed without gL, gD cell surface expression was unaffected, while gB cell surface expression was reduced. This further supports the hypothesis that gD does not interact with gH in the ER to an appreciable degree because gD is able to leave the ER and traffic to the cell surface unimpeded. By contrast, roughly a third of gB molecules appear to be held back from trafficking to the cell surface due to interactions with gH in the ER. Therefore, we propose that a portion of gH/gL and gB interact in the ER and traffic together to the cell surface in addition to some gB that traffics independently, whereas gD traffics to the cell surface independently and then joins the gH/gL-gB complex on the cell surface (**Fig. 9a**).

Both gD and gB interact with gH/gL through their ectodomains, TMDs, and CTDs, so it is interesting that gB but not gD interacted with gH in the ER. How is gD able to bind to gH/gL on the cell surface but avoid binding to gH in the ER? It is possible that the gD-gH interaction is more dynamic than the gH-gB interaction, with more frequent dissociation, allowing gD to gradually escape the ER without gH/gL. Another possibility is that the conformation of gH is different in the absence of gL than when it is complexed with gL, which has been suggested in several studies [58, 60, 68], causing decreased gD-gH interaction relative to gD-gH/gL interactions.

### New HSV-1 fusion pathway model

Collectively, we postulate that gD, gH/gL, and gB form a stable gD-gH/gL-gB complex, with gH/gL positioned in between gD and gB. A preassembled gD-gH/gL-gB complex in this orientation would enable efficient transmission of an activating signal from gD-receptor interaction to gH/gL to gB to trigger its fusogenic refolding (**Fig. 9b**). We propose that this signal transduction occurs by the following “conformational cascade” model. Upon receptor binding to HSV-1 gD, gD undergoes a conformational change [44] that activates the already bound gH/gL ectodomain. This causes a conformational change in the gH/gL ectodomain [35, 42] that is transmitted through the gH TMD and activates the gH CT to alter its interaction with the gB CTD [15, 69, 70]. Ultimately, this triggers gB to refold and cause fusion (**Fig. 9b**) [15]. Hence, if any of the HSV-1 gH domains are replaced, the signaling sequence from receptor to gB is disrupted and fusion cannot occur.

### Open questions

Here, we identify stable gD-gH/gL, gH/gL-gB, and gD-gB interactions, suggesting that pairs of these interacting partners exist in complexes with one another. However, it is still undetermined whether these pairs exist independently or whether they form the gD-gH/gL-gB complex. Our fusion pathway model (**Fig. 9b**) favors this latter possibility, which would position the glycoproteins ideally for rapid signal transduction upon receptor binding. It is yet unclear how these complexes change during fusion. The stable interactions observed here for each interaction pair suggest that the complexes do not associate or dissociate during fusion, but may, instead, change their conformation to fuse membranes. Ultimately, it is yet unclear how these glycoproteins interact on the virions. How glycoproteins are distributed on the HSV-1 surface and how they interact before and throughout the fusion process are important questions that beg in-depth investigation.

## MATERIALS AND METHODS

### Cells and plasmids

CHO cells [71] were gifts from J. M. Coffin and were grown in Ham’s F-12 medium with 10% fetal bovine serum (FBS), 100 IU penicillin, and 100 µg/ml streptomycin at 37° C and 5% CO2, except when noted otherwise. Plasmids pPEP98, pPEP99, pPEP100, and pPEP101 encode the full-length HSV-1 (strain KOS) gB, gD, gH, and gL genes, respectively, in a pCAGGS vector. These plasmids were gifts from P. G. Spear (Northwestern U.) [72]. Plasmids RLuc1-7 and RLuc8-11 (encoding the *Renilla* split luciferase genes) and pBG38 (encoding the nectin-1 gene) were gifts from G. H. Cohen and R. J. Eisenberg (U. Pennsylvania) [57, 73]. Plasmids pJLS11 (gB868) was generated previously [51]. Plasmids for the NanoBiT interaction assay, including PRKACA-Sm and PRKAR2A-Lg positive control plasmids, Halo-Sm negative control plasmid, and Sm- and Lg-BiT plasmids for tagging proteins of interest were purchased from Promega (Madison, WI) [48]. All plasmids from Promega contained an HSV-TK promoter. Plasmids p85 and p25 encode the full-length EBV gH and EBV gL genes in a pCAGGS vector, respectively, and were gifts from R. M. Longnecker (Northwestern U.).

### Cloning

The cloning strategies and primers used to generate the constructs used in this paper are detailed in the supplemental information section.

### Western blotting

Total cellular expression of NanoBiT constructs was tested using Western blotting. CHO cells were seeded at 2.5×10^5^ cells per well in 6-well plates. The next day, DNA constructs of interest were transfected. On day 3, cells were treated with RIPA buffer and a protease inhibitor and collected and spun down. The supernatants were mixed with SDS-PAGE loading dye and heated for 5 minutes at 95° C. Samples were separated by electrophoresis, transferred onto nitrocellulose membranes, and blocked. Strips of membranes were incubated with the appropriate primary antibody overnight at 4 °C. On day 4, membranes were incubated with fluorescent secondary antibodies for 1 hr at room temperature. Membranes were imaged using a LI-COR Odyssey imager. More detailed methods are included in the supplemental information section.

### Flow cytometry

Cell surface expression of gB, gH, and gD constructs were evaluated using flow cytometry. CHO cells were seeded at 2.5×10^5^ cells per well in 6-well plates. The next day, each well was transfected with DNA constructs of interest. On day 3, cells were collected and incubated for 1 hr on ice with appropriate primary antibodies. Cells were washed and incubated for 1 hr on ice in the dark with secondary antibodies, and washed again. The fluorescence of the cells was determined by flow cytometry. More detailed methods are included in the supplemental information section.

### NanoBiT interaction assay

Interactions between glycoproteins were measured using the NanoBiT interaction assay [48]. CHO cells were seeded into 6-well plates at 2.5×10^5^ cells per well for effector cells and 6-well plates at 2.5×10^5^ cells per well for target cells. The next day, effector cells were transfected with DNA of the two interacting partners and any remaining HSV-1 proteins required for fusion. Target cells was transfected with nectin-1 (pBG38) or pCAGGS. Four hours later, the effector cells were collected and seeded into 3 wells per condition of a 96-well plate. On day 3, the media of the effector cells was replaced with 40 µl per well of fusion medium (Ham’s F12 with 10% FBS, Penicillin/Streptomycin, 50 mM HEPES) with 1:50 Endurazine luciferase substrate (Promega) added. Cells were placed in a BioTek plate reader. Luminescence measurements were taken every 2 minutes for 1 hr at 37° C. Meanwhile, target cells were collected and resuspended in fusion medium. 40 µl of target cells were added to each well of effector cells. Luminescence measurements were taken every 2 minutes for 7.5 or 8 hrs. More detailed methods are included in the supplemental information section.

### Cell-cell fusion assay

Cell-cell fusion of gB, gH, and gD constructs was tested using a split-luciferase assay [57]. CHO cells were seeded into 3 wells per condition of a 96-well plate at 5×10^4^ cells per well for effector cells and 6-well plates at 2×10^5^ cells per well for target cells. The next day, effector cells were transfected with DNA of constructs of interest and part of a split luciferase (RLuc1-7). Each well of target cells was transfected with the complementary part of the split luciferase (RLuc8-11) and nectin-1. On day 3, the media of the effector cells was replaced with 40 µl per well of fusion medium with 1:500 Enduren luciferase substrate (Promega) added. Cells were incubated for 1 hr at 37° C. In the meantime, target cells were collected and resuspended in fusion medium. 40 µl of target cells were added to each well of effector cells. The plate was immediately placed in a BioTek plate reader. Luminescence measurements were taken every 2 minutes for 2 hr followed by measurements every hour for 6 hours. More detailed methods are included in the supplemental information section.

### Statistics

Statistical analysis was done for each experiment using GraphPad PRISM 9 software. An unpaired t-test with Welch’s correction was used to compare conditions to each other as shown.

## ACKNOWLEDGEMENTS

We thank Erin Sanders for help with cloning, flow cytometry, and fusion experiments. We thank Stephen Kwok and Allen Parmelee at the Tufts Flow Cytometry Core and Adam Hilterbrand for flow cytometry training. We thank Bing Dai for advice regarding Western blotting. We thank Scott Messenger (Promega) for technical advice regarding the NanoBiT interaction assay. We thank Roselyn J. Eisenberg and Gary H. Cohen (University of Pennsylvania) for the gift of plasmids and antibodies. We thank Richard Longnecker (Northwestern University) for the gift of plasmids and antibodies. We thank Patricia Spear (Northwestern University) for the gift of plasmids. We thank Andrew McGuire (Fred Hutchinson Cancer Research Center) for the gift of antibodies. We thank John Coffin, Marta Gaglia, Karl Munger, and Ellen White for helpful discussions. Research reported in this publication was supported by the National Institutes of Health under Award Numbers T32GM731042 (Z.P.), T32GM731043 (Z.P.), F30AI161795 (Z.P.), and 1R01AI164698 (E.E.H.), and by a Faculty Scholar grant 55108533 from Howard Hughes Medical Institute (E.E.H.). The content is solely the responsibility of the authors and does not necessarily represent the official views of the National Institutes of Health.

## AUTHOR CONTRIBUTIONS

Z.P. designed the experiments, cloned the constructs, conducted the experiments, analyzed the data, generated hypotheses, generated models, and wrote the manuscript. A.R.V. cloned constructs, conducted experiments, analyzed the data, and generated hypotheses. E.E.H. designed experiments, analyzed the data, generated hypotheses, generated models, and wrote the manuscript.

## Notes

### Competing Interest Statement

The authors have declared no competing interest.

### Summary of Updates

The typo in the last name of the middle author has been corrected.

## REFERENCES

1. Harrison, S.C., Viral membrane fusion. Nat Struct Mol Biol, 2008. 15(7): p. 690–8.

2. Connolly, S.A., T.S. Jardetzky, and R. Longnecker, The structural basis of herpesvirus entry. Nat Rev Microbiol, 2021. 19(2): p. 110–121.

3. Cohen, J.I., Herpesvirus latency. J Clin Invest, 2020. 130(7): p. 3361–3369.

4. Renner, D.W. and M.L. Szpara, Impacts of Genome-Wide Analyses on Our Understanding of Human Herpesvirus Diversity and Evolution. J Virol, 2018. 92(1).

5. Looker, K.J., et al., Global and Regional Estimates of Prevalent and Incident Herpes Simplex Virus Type 1 Infections in 2012. PLoS One, 2015. 10(10): p. e0140765.

6. Arduino, P.G. and S.R. Porter, Herpes Simplex Virus Type 1 infection: overview on relevant clinico-pathological features. J Oral Pathol Med, 2008. 37(2): p. 107–21.

7. Steiner, I. and F. Benninger, Update on Herpes Virus Infections of the Nervous System. Current Neurology and Neuroscience Reports, 2013. 13(12): p. 414.

8. Sauerbrei, A. and P. Wutzler, Herpes simplex and varicella-zoster virus infections during pregnancy: current concepts of prevention, diagnosis and therapy. Part 1: herpes simplex virus infections. Med Microbiol Immunol, 2007. 196(2): p. 89–94.

9. Whitley, R., D.W. Kimberlin, and C.G. Prober, Pathogenesis and disease, in Human Herpesviruses: Biology, Therapy, and Immunoprophylaxis, A. Arvin, et al., Editors. 2007, Cambridge University Press Copyright © Cambridge University Press 2007.: Cambridge.

10. Wang, L., et al., Construction, identification, and immunogenic assessments of an HSV-1 mutant vaccine with a UL18 deletion. Acta Virol, 2018. 62(2): p. 164–171.

11. Bernstein, D.I., et al., The HSV-1 live attenuated VC2 vaccine provides protection against HSV-2 genital infection in the guinea pig model of genital herpes. Vaccine, 2019. 37(1): p. 61–68.

12. Vollmer, B., et al., The prefusion structure of herpes simplex virus glycoprotein B. Sci Adv, 2020. 6(39).

13. Liu, Y., et al., Prefusion structure of human cytomegalovirus glycoprotein B and structural basis for membrane fusion. Sci Adv, 2021. 7(10).

14. Heldwein, E.E., et al., Crystal structure of glycoprotein B from herpes simplex virus 1. Science, 2006. 313(5784): p. 217–220.

15. Cooper, R.S., et al., Structural basis for membrane anchoring and fusion regulation of the herpes simplex virus fusogen gB. Nat Struct Mol Biol, 2018. 25(5): p. 416–424.

16. Burke, H.G. and E.E. Heldwein, Crystal Structure of the Human Cytomegalovirus Glycoprotein B. PLoS Pathog, 2015. 11(10): p. e1005227.

17. Vallbracht, M., et al., Structure-Function Dissection of Pseudorabies Virus Glycoprotein B Fusion Loops. J Virol, 2018. 92(1).

18. Backovic, M., R. Longnecker, and T.S. Jardetzky, Structure of a trimeric variant of the Epstein-Barr virus glycoprotein B. Proc Natl Acad Sci U S A, 2009. 106(8): p. 2880–5.

19. Chen, J., et al., Ephrin Receptor A4 is a New Kaposi’s Sarcoma-Associated Herpesvirus Virus Entry Receptor. mBio, 2019. 10(1).

20. Hahn, A.S., et al., The ephrin receptor tyrosine kinase A2 is a cellular receptor for Kaposi’s sarcoma–associated herpesvirus. Nat Med, 2012. 18(6): p. 961–6.

21. Yang, E., A.M. Arvin, and S.L. Oliver, Role for the αV Integrin Subunit in Varicella-Zoster Virus-Mediated Fusion and Infection. J Virol, 2016. 90(16): p. 7567–78.

22. Chen, J., et al., Ephrin receptor A2 is a functional entry receptor for Epstein-Barr virus. Nat Microbiol, 2018. 3(2): p. 172–180.

23. Gonzalez-Del Pino, G.L. and E.E. Heldwein, Well Put Together—A Guide to Accessorizing with the Herpesvirus gH/gL Complexes. Viruses, 2022. 14(2): p. 296.

24. Carfi, A., et al., Herpes simplex virus glycoprotein D bound to the human receptor HveA. Mol Cell, 2001. 8(1): p. 169–79.

25. Di Giovine, P., et al., Structure of herpes simplex virus glycoprotein D bound to the human receptor nectin-1. PLoS Pathog, 2011. 7(9): p. e1002277.

26. Spear, P.G., Herpes simplex virus: receptors and ligands for cell entry. Cell Microbiol, 2004. 6(5): p. 401–10.

27. Martinez-Martin, N., et al., An Unbiased Screen for Human Cytomegalovirus Identifies Neuropilin-2 as a Central Viral Receptor. Cell, 2018. 174(5): p. 1158–1171.e19.

28. Kabanova, A., et al., Platelet-derived growth factor-α receptor is the cellular receptor for human cytomegalovirus gHgLgO trimer. Nat Microbiol, 2016. 1(8): p. 16082.

29. Wu, Y., et al., Human cytomegalovirus glycoprotein complex gH/gL/gO uses PDGFR-α as a key for entry. PLoS Pathog, 2017. 13(4): p. e1006281.

30. Mullen, M.M., et al., Structure of the Epstein-Barr virus gp42 protein bound to the MHC class II receptor HLA-DR1. Mol Cell, 2002. 9(2): p. 375–85.

31. Sathiyamoorthy, K., et al., Structural basis for Epstein-Barr virus host cell tropism mediated by gp42 and gHgL entry glycoproteins. Nat Commun, 2016. 7: p. 13557.

32. Chandramouli, S., et al., Structural basis for potent antibody-mediated neutralization of human cytomegalovirus. Sci Immunol, 2017. 2(12).

33. Ciferri, C., et al., Antigenic Characterization of the HCMV gH/gL/gO and Pentamer Cell Entry Complexes Reveals Binding Sites for Potently Neutralizing Human Antibodies. PLoS Pathog, 2015. 11(10): p. e1005230.

34. Si, Z., et al., Different functional states of fusion protein gB revealed on human cytomegalovirus by cryo electron tomography with Volta phase plate. PLoS Pathog, 2018. 14(12): p. e1007452.

35. Cairns, T.M., et al., Localization of the Interaction Site of Herpes Simplex Virus Glycoprotein D (gD) on the Membrane Fusion Regulator, gH/gL. J Virol, 2020. 94(20).

36. Cairns, T.M., et al., Surface Plasmon Resonance Reveals Direct Binding of Herpes Simplex Virus Glycoproteins gH/gL to gD and Locates a gH/gL Binding Site on gD. J Virol, 2019. 93(15).

37. Atanasiu, D., et al., Bimolecular complementation reveals that glycoproteins gB and gH/gL of herpes simplex virus interact with each other during cell fusion. Proc Natl Acad Sci U S A, 2007. 104(47): p. 18718–23.

38. Avitabile, E., C. Forghieri, and G. Campadelli-Fiume, Complexes between herpes simplex virus glycoproteins gD, gB, and gH detected in cells by complementation of split enhanced green fluorescent protein. J Virol, 2007. 81(20): p. 11532–7.

39. Avitabile, E., C. Forghieri, and G. Campadelli-Fiume, Cross talk among the glycoproteins involved in herpes simplex virus entry and fusion: the interaction between gB and gH/gL does not necessarily require gD. J Virol, 2009. 83(20): p. 10752–60.

40. Atanasiu, D., et al., Cascade of events governing cell-cell fusion induced by herpes simplex virus glycoproteins gD, gH/gL, and gB. J Virol, 2010. 84(23): p. 12292–9.

41. Cocchi, F., et al., The soluble ectodomain of herpes simplex virus gD contains a membrane-proximal pro-fusion domain and suffices to mediate virus entry. Proc Natl Acad Sci U S A, 2004. 101(19): p. 7445–50.

42. Atanasiu, D., et al., Regulation of herpes simplex virus gB-induced cell-cell fusion by mutant forms of gH/gL in the absence of gD and cellular receptors. mBio, 2013. 4(2).

43. Heldwein, E.E. and C. Krummenacher, Entry of herpesviruses into mammalian cells. Cell Mol Life Sci, 2008. 65(11): p. 1653–68.

44. Krummenacher, C., et al., Structure of unliganded HSV gD reveals a mechanism for receptor-mediated activation of virus entry. Embo j, 2005. 24(23): p. 4144–53.

45. Zhang, N., et al., Binding of herpes simplex virus glycoprotein D to nectin-1 exploits host cell adhesion. Nat Commun, 2011. 2: p. 577.

46. Chowdary, T.K., et al., Crystal structure of the conserved herpesvirus fusion regulator complex gH-gL. Nat Struct Mol Biol, 2010. 17(7): p. 882–8.

47. Romei, M.G. and S.G. Boxer, Split Green Fluorescent Proteins: Scope, Limitations, and Outlook. Annu Rev Biophys, 2019. 48: p. 19–44.

48. Dixon, A.S., et al., NanoLuc Complementation Reporter Optimized for Accurate Measurement of Protein Interactions in Cells. ACS Chem Biol, 2016. 11(2): p. 400–8.

49. Promega NanoBiT Protein:Protein Interaction System Technical Manual. http://www.promega.com, accessed on 3/6/22, 2021.

50. England, C.G., H. Luo, and W. Cai, HaloTag technology: a versatile platform for biomedical applications. Bioconjug Chem, 2015. 26(6): p. 975–86.

51. Silverman, J.L., et al., Membrane requirement for folding of the herpes simplex virus 1 gB cytodomain suggests a unique mechanism of fusion regulation. J Virol, 2012. 86(15): p. 8171–84.

52. Zeev-Ben-Mordehai, T., et al., Extracellular vesicles: a platform for the structure determination of membrane proteins by Cryo-EM. Structure, 2014. 22(11): p. 1687–92.

53. Zeev-Ben-Mordehai, T., et al., Two distinct trimeric conformations of natively membrane-anchored full-length herpes simplex virus 1 glycoprotein B. Proc Natl Acad Sci U S A, 2016. 113(15): p. 4176–81.

54. Maurer, U.E., et al., The structure of herpesvirus fusion glycoprotein B-bilayer complex reveals the protein-membrane and lateral protein-protein interaction. Structure, 2013. 21(8): p. 1396–405.

55. Lee, S.K., T. Compton, and R. Longnecker, Failure to complement infectivity of EBV and HSV-1 glycoprotein B (gB) deletion mutants with gBs from different human herpesvirus subfamilies. Virology, 1997. 237(1): p. 170–81.

56. Vanarsdall, A.L., et al., Human Cytomegalovirus gH/gL Forms a Stable Complex with the Fusion Protein gB in Virions. PLoS Pathog, 2016. 12(4): p. e1005564.

57. Saw, W.T., et al., Using a split luciferase assay (SLA) to measure the kinetics of cell-cell fusion mediated by herpes simplex virus glycoproteins. Methods, 2015. 90: p. 68–75.

58. Hutchinson, L., et al., A novel herpes simplex virus glycoprotein, gL, forms a complex with glycoprotein H (gH) and affects normal folding and surface expression of gH. J Virol, 1992. 66(4): p. 2240–50.

59. Cairns, T.M., et al., N-terminal mutants of herpes simplex virus type 2 gH are transported without gL but require gL for function. J Virol, 2007. 81(10): p. 5102–11.

60. Roberts, S.R., et al., Analysis of the intracellular maturation of the herpes simplex virus type 1 glycoprotein gH in infected and transfected cells. Virology, 1991. 184(2): p. 609–24.

61. Atanasiu, D., et al., Using Split Luciferase Assay and anti-HSV Glycoprotein Monoclonal Antibodies to Predict a Functional Binding Site Between gD and gH/gL. J Virol, 2021. 95(8).

62. Gianni, T., R. Massaro, and G. Campadelli-Fiume, Dissociation of HSV gL from gH by alphavbeta6-or alphavbeta8-integrin promotes gH activation and virus entry. Proc Natl Acad Sci U S A, 2015. 112(29): p. E3901–10.

63. Gianni, T., et al., αvβ6-and αvβ8-integrins serve as interchangeable receptors for HSV gH/gL to promote endocytosis and activation of membrane fusion. PLoS Pathog, 2013. 9(12): p. e1003806.

64. Stampfer, S.D. and E.E. Heldwein, Stuck in the middle: structural insights into the role of the gH/gL heterodimer in herpesvirus entry. Curr Opin Virol, 2013. 3(1): p. 13–9.

65. Hollinshead, M., et al., Endocytic tubules regulated by Rab GTPases 5 and 11 are used for envelopment of herpes simplex virus. Embo j, 2012. 31(21): p. 4204–20.

66. Albecka, A., et al., HSV-1 Glycoproteins Are Delivered to Virus Assembly Sites Through Dynamin-Dependent Endocytosis. Traffic, 2016. 17(1): p. 21–39.

67. Hogue, I.B., J. Scherer, and L.W. Enquist, Exocytosis of Alphaherpesvirus Virions, Light Particles, and Glycoproteins Uses Constitutive Secretory Mechanisms. mBio, 2016. 7(3).

68. Gompels, U.A. and A.C. Minson, Antigenic properties and cellular localization of herpes simplex virus glycoprotein H synthesized in a mammalian cell expression system. J Virol, 1989. 63(11): p. 4744–55.

69. Rogalin, H.B. and E.E. Heldwein, Interplay between the Herpes Simplex Virus 1 gB Cytodomain and the gH Cytotail during Cell-Cell Fusion. J Virol, 2015. 89(24): p. 12262–72.

70. Pataki, Z., E.K. Sanders, and E.E. Heldwein, A surface pocket in the cytoplasmic domain of the herpes simplex virus fusogen gB controls membrane fusion. bioRxiv, 2022: p. 2022.03.14.484201.

71. Puck, T.T., S.J. Cieciura, and A. Robinson, Genetics of somatic mammalian cells. III. Long-term cultivation of euploid cells from human and animal subjects. J Exp Med, 1958. 108(6): p. 945–56.

72. Pertel, P.E., et al., Cell fusion induced by herpes simplex virus glycoproteins gB, gD, and gH-gL requires a gD receptor but not necessarily heparan sulfate. Virology, 2001. 279(1): p. 313–24.

73. Krummenacher, C., et al., Effects of herpes simplex virus on structure and function of nectin-1/HveC. J Virol, 2002. 76(5): p. 2424–33.

74. Gibson, D.G., et al., Enzymatic assembly of DNA molecules up to several hundred kilobases. Nat Methods, 2009. 6(5): p. 343–5.

75. Arii, J., et al., Entry of herpes simplex virus 1 and other alphaherpesviruses via the paired immunoglobulin-like type 2 receptor alpha. J Virol, 2009. 83(9): p. 4520–7.

