## Supplemental Materials and Methods and Supplemental Table 1 for "Herpes simplex virus 1 entry glycoproteins form stable complexes prior to and during membrane fusion"

**Cloning of HSV-1 gB constructs tagged with Sm-BiT.** To generate HSV-1 gB-Sm, pPEP98 was digested with XhoI and NheI to isolate the largest fragment of the gB gene for insertion into the NanoBiT plasmid. The remaining 3’ end of the gene missing from the XhoI-NheI fragment was PCR amplified using the forward primer P1 and the reverse primer P2. The Lg-BiT plasmid was amplified with the forward primer P3 and reverse primer P4 to generate overhangs with the gB gene for Gibson assembly. The Lg-BiT DNA and two fragments of gB DNA were assembled using Gibson assembly [74] with the Gibson Assembly Master Mix (New England Biolabs, Ipswich, MA) to generate gB-Lg HSV-TK. This plasmid was then digested at an XhoI and XbaI site to remove the Lg-BiT tag. The Sm-BiT tag was amplified from the Sm-BiT plasmid using forward primer P5 and reverse primer P6 to generate overhangs with the gB gene lacking the Lg-BiT tag. The two DNA fragments were assembled using Gibson assembly to generate gB-Sm HSV-TK. This plasmid was digested at a KpnI and HindIII site to remove the HSV-TK promoter. A higher expression CAG enhancer was PCR amplified from pPEP98 using the forward primer P7 and the reverse primer P8 to generate overhangs with the gB-Sm fragment lacking the HSV-TK promoter. The new CAG promoter was inserted into the gB-Sm fragment using Gibson assembly to generate the gB-Sm plasmid used in this work.

To generate gB-Sm H516P, gB-Sm was digested with AgeI and PmlI to generate the backbone for Gibson assembly. The forward primer P9 and reverse primer P10 were used to generate the first DNA fragment, and the forward primer P11 and reverse primer P12 were used to generate the second DNA fragment for insertion. The two fragments were assembled with the AgeI-PmlI plasmid backbone using Gibson assembly to generate gB-Sm H516P. The resulting plasmids were pZP42 (gB-Sm) and pZP70 (gB-Sm H516P). All primers are listed in Supplemental table 1.

**Cloning of gH constructs tagged with Lg-BiT.** To generate HSV-1 gH-Lg, pPEP100 was digested with SacI and MfeI to isolate the largest fragment of the gH gene for insertion into the NanoBiT plasmid. The remaining 3’ end of the gene missing from the SacI-MfeI fragment was PCR amplified using the forward primer P13 and reverse primer P14. The Lg-BiT plasmid was amplified with the forward primer P15 and reverse primer P16 to generate overhangs with the gH gene for Gibson assembly. The Lg-BiT DNA and two gH fragments were assembled using Gibson assembly to generate gH-Lg HSV-TK. The HSV-TK promoter was swapped for the CAG enhancer using the same strategy as described above to generate the HSV-1 gH-Lg plasmid used in this work.

To generate gH-Lg with the HSV-1 TMD replaced with that of EBV (gH-Lg TMD_EBV_), pPEP100 was digested with SacI and BspEI to generate a backbone for Gibson assembly. pPEP100 was amplified with the forward primer P17 and reverse primer P18 to obtain the HSV-1 ectodomain fragment. The EBV gH plasmid was amplified with the forward primer P19 and the reverse primer P20 to obtain EBV TMD and HSV-1 CT fragment. The two fragments were inserted into the cut plasmid backbone by Gibson assembly. This plasmid was used as a PCR template with forward primer P21 and reverse primer P22 to generate the insert for assembly. The gH-Lg plasmid with the CAG promoter was digested with StuI and XhoI to obtain the vector backbone for Gibson assembly to generate gH-Lg TMD_EBV_.

To generate HSV-1 gH-Lg with a scrambled CT (gH-Lg CT_SCR_), HSV-1 gH-Lg was amplified using the forward primer P23 and the reverse primer P24. To extend this DNA product, it was amplified using the forward primer P23 and the reverse primer P25. This product was inserted into the gH-Lg backbone digested with StuI and XhoI using Gibson assembly to obtain gH-Lg CT_SCR_.

To generate HSV-1 gH-Lg with the ectodomain replaced with EBV (gH-Lg ECTO_EBV_), the gH-Lg plasmid was digested with PacI and XhoI to create the backbone for the assembly. The EBV gH plasmid was amplified with the forward primer P26 and reverse primer P27 to generate the EBV ectodomain. pPEP100 was amplified with the forward primer P28 and reverse primer P22 to generate the HSV-1 TMD and CT. The two DNA fragments were inserted into the vector backbone by Gibson assembly to create gH-Lg ECTO_EBV_.

To generate HSV-1 gH-Lg with the TMD replaced with EBV and the CT scrambled (gH-Lg TMD_EBV_-CT_SCR_), gH-Lg TMD_EBV_ was amplified using the forward primer P29 and reverse primer P30. This product was further extended using the forward primer P29 and reverse primer P25 and then inserted into the gH-Lg backbone digested by StuI and XhoI by Gibson assembly to generate gH-Lg TMD_EBV_-CT_SCR_.

To generate HSV-1 gH-Lg with the ectodomain and TMD replaced with EBV (gH-Lg ECTO_EBV_-TMD_EBV_), gH-Lg ECTO_EBV_ was digested with BlpI and XhoI to generate the vector backbone for the assembly. The EBV gH plasmid was amplified using forward primer P31 and reverse primer P20 to obtain the EBV ectodomain and TMD. This product was extended using forward primer P31 and reverse primer P22 and then inserted into the vector backbone by Gibson assembly to create gH-Lg ECTO_EBV_-TMD_EBV_.

To generate HSV-1 gH-Lg with the ectodomain replaced with EBV and the CT scrambled (gH-Lg ECTO_EBV_-CT_SCR_), gH-Lg ECTO_EBV_ was amplified using forward primer P31 and reverse primer P24. This product was extended using the forward primer P31 and the reverse primer P25 and then inserted into the gH-Lg ECTO_EBV_ backbone digested with BlpI and XhoI by Gibson assembly to create gH-Lg ECTO_EBV_-CT_SCR_.

To generate EBV gH-Lg, the EBV gH plasmid was amplified using forward primer P31 and reverse primer P32 to obtain the EBV gH gene with Gibson overhangs. This PCR product was inserted into the gH-Lg ECTO_EBV_ backbone digested with BlpI and XhoI to generate EBV gH-Lg by Gibson assembly. The resulting plasmids were pZP44 (gH-Lg), pZP71 (gH-Lg TMD_EBV_), pZP68 (gH-Lg CT_SCR_), pZP73 (gH-Lg ECTO_EBV_), pZP77 (gH-Lg TMD_EBV_-CT_SCR_), pZP75 (gH-Lg ECTO_EBV_-TMD_EBV_), pZP76 (gH-Lg ECTO_EBV_-CT_SCR_), pZP80 (EBV gH-Lg). All primers are listed in Supplemental table 1. EBV gH and gL plasmids were gifts from R. M. Longnecker (Northwestern U.).

**Cloning of HSV-1 gD constructs tagged with Sm- and Lg-BiT.** To generate HSV-1 gD-Sm, gB-Sm was digested with PacI and XhoI to generate the vector backbone for the assembly. pPEP99 was amplified using the forward primer P33 and reverse primer P34 to obtain the gD gene with Gibson overhangs. This PCR product was inserted into the vector backbone by Gibson assembly to generate gD-Sm. To generate HSV-1 gD-Lg, gD-Sm was digested with XhoI and XbaI to generate the vector backbone for the assembly. gH-Lg was PCR amplified using forward primer P35 and reverse primer P36 to obtain the insert for the assembly. The insert and backbone were joined using Gibson assembly to generate gD-Lg. The resulting plasmids were pZP78 (gD-Sm) and pZP79 (gD-Lg). All primers are listed in Supplemental table 1.

**Cloning of Halo-Sm construct.** The HSV-TK promoter of the Halo-Sm negative control plasmid was also replaced with a CAG enhancer, as follows. Halo-Sm HSV-TK was digested with NheI and BglII to remove the HSV-TK promoter. The CAG enhancer was PCR amplified from pPEP98 using the forward primer P37 and the reverse primer P38 to create overhangs with the Halo-Sm plasmid with the promoter cut out. To extend the 3’ overhangs further, this PCR product was amplified with the forward primer P37 and the reverse primer P39. This DNA fragment was assembled with the Halo-Sm plasmid lacking the promoter using Gibson assembly to generate the Halo-Sm plasmid used in this work (pZP49). All primers are listed in Supplemental table 1.

**Western blotting.** Total cellular expression of NanoBiT constructs was tested using Western blotting. CHO cells were seeded at 2.5x10^5^ cells per well in 6-well plates. The next day, 2 µg of gB construct (pPEP98 or gB-Sm construct) or gD construct (pPEP99 or gD-Sm or gD-Lg) or 1 µg each of gL (pPEP101) and gH constructs (pPEP100 or gH-Lg construct) were transfected using 4 µl of JetPrime (Polyplus, Illkirch-Graffenstaden, France) and 200 µl of JetPrime buffer per well. In the experiment where gH-Lg was transfected without gL, pCAGGS empty vector was transfected in the place of gL. 2 µg of pCAGGS was transfected as a negative control in all experiments. On day 3, a protease inhibitor (Sigma-Aldrich, St. Louis, MO) was added to RIPA buffer, which was added to the cells, and the cells were collected and spun down. The supernatants were mixed with SDS-PAGE loading dye and heated for 5 minutes at 95° C. Samples were loaded onto 4-15% precast polyacrylamide gels (Bio-Rad, Hercules, CA) and separated by electrophoresis. Samples were transferred onto nitrocellulose membranes and blocked with 5% milk in TBST. Strips of membranes were incubated with the appropriate primary antibody in 5% milk in TBST overnight at 4 °C. Rabbit anti-HSV-1 gB polyclonal antibody (pAb) R68 was used at 1:10,000, rabbit anti-HSV-1 gH pAb R137 was used at 1:5,000, rabbit anti-EBV gH/gL pAb R2267 was used at 1:1,000, rabbit anti-HSV-1 gD pAb R7 was used at 1:10,000, and rabbit anti-tubulin pAb was used at 1:1,000 against tubulin (9F3, Cell Signaling Technology, Danvers, MA) as a loading control. On day 4, membranes were washed and incubated with IRDye 800CW anti-rabbit fluorescent secondary antibodies (LI-COR Biosciences, Lincoln, NE) at 1:5,000 in 5% milk in TBST for 1 hr at room temperature. Membranes were washed and imaged using a LI-COR Odyssey imager. Band intensities were calculated using Image Studio. Band intensities were divided by the tubulin band intensity of the same well. Reported values are averages of three independent experiments. R68, R137, and R7 pAbs were gifts from G. H. Cohen and R. J. Eisenberg (U. Pennsylvania). R2267 pAb was a gift from R. M. Longnecker (Northwestern U.).

**Flow cytometry.** Cell surface expression of gB, gH, and gD constructs were evaluated using flow cytometry. CHO cells were seeded at 2.5x10^5^ cells per well in 6-well plates. The next day, each well was transfected with 2 µg gB (pPEP98 or gB-Sm construct) or gD (pPEP99 or gD-Sm or gD-Lg) or 1 µg each of gH (pPEP100 or gH-Lg construct) plus 1 µg gL (pPEP101) using 4 µl of JetPrime in 200 µl JetPrime buffer. Under conditions testing gH without gL, 1 µg of pCAGGS was transfected instead of gL. As a negative control, 2 µg of pCAGGS was transfected instead of viral genes. One well was left untransfected as a ‘mock’ condition. On day 3, cells were detached with 1 ml per well of Versene (Fisher Scientific, Waltham, MA) and collected using FACS media (PBS (Cytiva, Marlborough, MA) with 3% FBS). Cells were washed with FACS media twice. Cells were incubated for 1 hr on ice with 250 µl of 1:500 primary antibody rabbit anti-HSV-1 gB pAb R68 and mouse anti-HSV-1 gH/gL mAb LP11 or 1:1000 for rabbit anti-HSV-1 gH/gL pAb R137, human anti-EBV gH/gL mAb AMMO1 and mouse anti-HSV-1 gD mAb DL6 in FACS media. A pCAGGS negative control condition was also incubated with these antibodies in each experiment. Anti-c-myc rabbit (A14, Santa Cruz Biotechnology) or mouse (9E10, Santa Cruz Biotechnology, Dallas, TX) antibodies were used as a non-targeting negative control for the Mock condition. Cells were washed three times and incubated for 1 hr on ice in the dark with 250 µl 1:250 secondary FITC-conjugated anti-rabbit antibody (MPBio, Santa Ana, CA) for R68 conditions, 1:250 Alexa Fluor 488 anti-mouse antibody (Invitrogen, Waltham, MA) for LP11 conditions, 1:500 FITC-conjugated anti-rabbit antibody for R137 conditions, and 1:500 Alexa Fluor 488 anti-mouse antibody for DL6 conditions. AMMO1 is conjugated to Dylite-650, so no secondary antibody was needed. Cells were washed three times. The fluorescence of the cells was determined by flow cytometry. Gating of live cells was performed using forward scatter (FSC) and side scatter (SSC) using FlowJo software. gB+, gH+, and gD+ cells were gated using the pCAGGS condition as a negative control, using a cutoff of 5% gB+, gH+, or gD+ pCAGGS cells to capture the greater majority of true positives while minimizing false positives. Total cell surface expression of the transfected population was obtained by calculating the product of % gB+, gH+, or gD+ cells and the mean fluorescence intensity of the gB+, gH+, or gD+ cells. Total cell surface expression was then normalized to the WT gB, WT gH, WT gD, gB-Sm, or gH-Lg condition, expressed as a percentage. The values represent the average of three independent experiments. LP11 (monoclonal anti-HSV-1 gH/gL) was a gift from H. Browne (U. of Cambridge). AMMO1 (monoclonal anti-EBV gH/gL) was a gift from A. T. McGuire (Fred Hutchinson Cancer Research Center).

**NanoBiT interaction assay.** Interactions between glycoproteins were measured using the NanoBiT interaction assay [48]. CHO cells were seeded into 6-well plates at 2.5x10^5^ cells per well for effector cells and 6-well plates at 2.5x10^5^ cells per well for target cells. The next day, effector cells were transfected per well with 638 ng of one interacting partner (gB-Sm construct, gD-Sm construct, Halo-Sm, PRKACA, or untagged protein), 638 ng of the complementary interacting partner (gH-Lg construct, gD-Lg construct, PRKAR2A, or untagged protein), and 213 ng of any remaining HSV-1 proteins required for fusion, such as gL, or pCAGGS in the PRKACA/PRKAR2A condition, using 3.4 µl JetPrime in 200 µl JetPrime buffer. Whenever certain plasmids were omitted to test their effects on the interaction, pCAGGS was transfected in their place. CHO cells lack HSV-1 receptors, so no fusion can occur until receptor-bearing target cells are introduced [75]. Each well of target cells was transfected with 1 µg of the HSV-1 receptor nectin-1 (pBG38) or pCAGGS with 2 µl of JetPrime in 200 µl of JetPrime buffer. Four hours later, the effector cells were detached by incubating with 1 ml per well of Versene. Effector cells were collected, spun down, and resuspended in 500 µl of culture medium per well. 100 µl of effector cells were seeded into 3 wells per condition of a 96-well plate. On day 3, the tissue culture media was removed from the 96-well plate wells and replaced with 40 µl per well of fusion medium (Ham’s F12 with 10% FBS, Penicillin/Streptomycin, 50 mM HEPES) with 1:50 Endurazine luciferase substrate (Promega) added. The Endurazine concentration becomes 1:100 once the target cells are added. Cells were placed in a BioTek plate reader. Luminescence measurements were taken every 2 minutes for 1 hr at 37º C to measure interaction before fusion. Meanwhile, target cells were detached by incubating with 1 ml per well of Versene. Target cells were collected, spun down, and resuspended in 500 µl of fusion medium per well. 40 µl of target cells were added to each well of effector cells to trigger fusion. The plate was placed in a BioTek plate reader. Luminescence measurements were taken every 2 minutes for 7.5 or 8 hrs to measure interaction during fusion. Untagged proteins and Halo-Sm, which is an unrelated protein that is not known to interact with viral proteins, paired with a Lg-tagged protein of interest, served as negative controls. PRKACA/PRKAR2A are two tagged proteins known to interact, serving as a positive control. Halo-Sm and PRKACA/PRKAR2A conditions were included in every experiment to ensure the assay was working as expected. To generate interaction curves, luminescence values were averaged for the three wells in each condition and normalized to the maximum signal of the Halo-Sm negative control condition, expressed as a fold change over the negative control. To calculate the total interaction over the time course, the area under the interaction curves generated from raw values was calculated using GraphPad PRISM 9 software. The area under the curve (AUC) was then normalized to the AUC of the Halo-Sm condition, expressed as a fold change over the negative control. Alternatively, for comparisons between similar conditions, the AUC of experimental conditions was normalized to the AUC of a control condition, expressed as a percentage. Reported values are averages of three independent experiments.

**Cell-cell fusion assay.** Cell-cell fusion of gB, gH, and gD constructs was tested using a split-luciferase assay [57]. CHO cells were seeded into 3 wells per condition of a 96-well plate at 5x10^4^ cells per well for effector cells and 6-well plates at 2x10^5^ cells per well for target cells. The next day, effector cells were transfected per well with 125 ng each of Sm- or Lg-tagged interacting partners (gB, gH, or gD) or untagged control and 41.7 ng each of split luciferase (RLuc1-7) and remaining untagged viral proteins required for fusion, such as gL, using 0.75 µl JetPrime in 10 µl JetPrime buffer. For the pCAGGS negative control condition, 333 ng of pCAGGS empty vector was transfected instead of the gB, gH, gL, and gD. Each well of target cells was transfected with 1 µg of the complementary part of the split luciferase (RLuc8-11) and 1 µg of the HSV-1 receptor nectin-1 (pBG38) with 4 µl of JetPrime in 200 µl of JetPrime buffer. On day 3, the tissue culture media in the 96-well plate wells was replaced with 40 µl per well of fusion medium with 1:500 Enduren luciferase substrate (Promega) added. The Enduren concentration becomes 1:1,000 after target cells are added. Cells were incubated for 1 hr at 37º C. In the meantime, target cells were detached by incubating with 1 ml per well of Versene. The target cells were collected, spun down, and resuspended in 500 µl of fusion medium per well. 40 µl of target cells were added to each well of effector cells. The plate was immediately placed in a BioTek plate reader. Luminescence measurements were taken every 2 minutes for 2 hr followed by measurements every hour for 6 hours. A condition containing gB868 instead of untagged or tagged gB was always included as a hyperfusogenic positive control to make sure that the assay is working as expected. The average hyperfusogenic positive control signal was always higher than that of the WT gB condition in all experiments. The NanoBiT tags cannot utilize the Enduren fusion assay luciferase substrate to produce luminescence. Nevertheless, controls were included for each condition in which target cells lacking nectin-1 receptor were added instead of target cells expressing nectin-1 to evaluate any background luminescence caused by the NanoBiT tags in the absence of fusion. The luminescence in these conditions was at the level of the pCAGGS negative control condition. Luminescence values were averaged for the three wells in each condition and normalized to the signal at 8 hr in the condition with untagged WT proteins, expressed as a percentage of WT. Reported values are averages of three independent experiments.

**Supplemental table 1. Primers used for cloning.**

| **Primer** | **Sequence (5’-3’)** | **Primer** | **Sequence (5’-3’)** |
| --- | --- | --- | --- |
| P1 | GCGACTTTGACGAGGCCAAGCTAGCCGAGGCCAGGGAGATGATACGGTACATG | P21 | GCTCAGGGGAATTCTGGTTAATTAACGGTACCCGGGTCCCCCATGGGGAATG |
| P2 | GCCACCACCGCTCGAGAGGTCGTCCTCGTCGGCGTCAC | P22 | ACCTCCGCTCCCGCCACCACTCGAGCCTTCGCGTCTCCAAAAAAACGGGACACTTGTCCG |
| P3 | ACGAGGACGACCTCTCGAGCGGTGGTGGC | P23 | GCCTTCGTCCCTGAGGCCTCACATCGGTGCGGGGGGCAGTCT |
| P4 | TGCATGGCAGATCGCCTAGCTTAATTAACCAGAATTCCCCTGAGCTCC | P24 | ACTTGTGACTCTAAACTTGCGCGGCCAGAGAAATTCCCGTAGGATGCCAGCCAGGGCGGC |
| P5 | GACGCCGACGAGGACGACCTCTCGAGCGGTGGTGGC | P25 | ACCTCCGCTCCCGCCACCACTCGAGCCAACACTTGTGACTCTAAACTTGCGCGGCCAGAG |
| P6 | TCAAGGGCATCGGTCGACG | P26 | GCTCAGGGGAATTCTGGTTAATTAACGGTACCCGGGTCCCCCATGCAGTTG |
| P7 | AACATTTCTCTGGCCTAACTGGCCGGTACCTGAGTCTCGTTACATAACTTACGGTAAATG | P27 | GGCGGCCAGAAACCCTGCTCTTTCTTCATACAGGCCCGCAATTTCCATGACAGT |
| P8 | CAACAGTACCGGATTGCCAAGCTTCATGGTAATAGCGATGACTAATACG | P28 | AGAAAGAGCAGGGTTTCTGGCCGCCTCTGCGCT |
| P9 | ACCACCGACCTCAAGTACAACC | P29 | AAGTCCTGGCCCAGCAGACC |
| P10 | GTTGACAGGGTGCTGTATGTGGTTG | P30 | ACTCTAAACTTGCGCGGCCAGAGAAATTCCCGAACCAGAAAGATACCCAGAGCAAAAGC |
| P11 | ACATACAGCACCCTGTCAACGATATGTT | P31 | GACCTCACACGAGACAAGCTGC |
| P12 | TTGGACGATCACGTTGTCCGC | P32 | CGCTCCCGCCACCACTCGAGCCAAGGAAAAACATAACAATCTTGTGAACCAGAAAGATAC |
| P13 | CACGCAGCCCGTGGCCGCAATTGCGCCCGGGTTTCTGGCCGC | P33 | GCTCAGGGGAATTCTGGTTAATTAAGCTAGGCGATCTGCCATGGGGGGGGCTGCCGCCAG |
| P14 | GCCACCACTCGAGCCTTCGCGTCTCCAAAAAAACGG | P34 | TCCGCTCCCGCCACCACCGCTCGAGTAAAACAAGGGCTGGTGCGAGGACGGCTG |
| P15 | GCGAAGGCTCGAGTGGTGGCGGGAGCGGAGGTGGAGGGTC | P35 | CGCACCAGCCCTTGTTTTACGGCTCGAGTGGTGGCGGGAGCGGAGG |
| P16 | CATGGGGGACCCGGGTACCGTTAATTAACCAGAATTCCCCTGAGCTCCCACTTAGGCG | P36 | TATCTTATCATGTCTGCTCGAAGCGGCCGGCCGCC |
| P17 | ATCATTTTGGCAAAGAATTCGAGCTCGGTACC | P37 | GGCGTAGAGGATCGAGATCTGTACCTGAGTCTCGTTACATAACTTACGGTAAATGG |
| P18 | AACAACGTGGGGCGCAATTGCGGCCAC | P38 | GGTGGCTTTACCAACAGTACCGGATTGCCAAGCTTCATGGTAATAGCGATGACTAATACG |
| P19 | CAATTGCGCCCCACGTTGTTTTGGCAATAATCC | P39 | CTGCCATGGCGATCGCTAGCGGTGGCTTTACCAACAGTAC |
| P20 | CCAAAAAAACGGGACACTTGTCCGGAGAACCTTAACCAGAAAGATACCCAGAGC |  |  |
